# Multi-omics analysis identifies symbionts and pathogens of blacklegged ticks (*Ixodes scapularis*) from a Lyme disease hotspot in southeastern Ontario, Canada

**DOI:** 10.1101/2022.11.09.515820

**Authors:** Amber R. Paulson, Stephen C. Lougheed, David Huang, Robert I. Colautti

## Abstract

Ticks in the family *Ixodidae* are important vectors of zoonoses including Lyme disease (LD), which is caused by spirochete bacteria from the *Borreliella* (*Borrelia*) *burgdorferi* sensu lato (*Bbsl*) complex. The blacklegged tick (*Ixodes scapularis*) continues to expand across Canada, creating hotspots of elevated LD risk at the leading edge of its expansion range. Current efforts to understand the risk of pathogen transmission associated with *I. scapularis* in Canada focus primarily on targeted screens, while variation in the tick microbiome remains poorly understood. Using multi-omics consisting of 16S metabarcoding and ribosome-depleted, whole-shotgun RNA transcriptome sequencing, we examined the microbial communities associated with adult *I. scapularis* (N = 32), sampled from four tissue types (whole tick, salivary glands, midgut, and viscera) and three geographical locations within a LD hotspot near Kingston, Ontario, Canada. The communities consisted of both endosymbiotic and known or potentially pathogenic microbes, including RNA viruses, bacteria, and a *Babesia* sp. intracellular parasite. We show that β-diversity is significantly higher between individual tick salivary gland and midgut bacterial communities, compared to whole ticks; while linear discriminant analysis (LDA) effect size (LEfSe) determined that the three potentially pathogenic bacteria detected by V4 16S rDNA sequencing also differed among dissected tissues only, including a *Borrelia* from the *Bbsl* complex, *Borrelia miyamotoi*, and *Anaplasma phagocytophilum*. Importantly, we find co-infection of *I. scapularis* by multiple microbes, in contrast to diagnostic protocols for LD, which typically focus on infection from a single pathogen of interest (*B. burgdorferi* sensu stricto).

**IMPORTANCE:** A vector of human health concern, blacklegged ticks, *Ixodes scapularis*, transmit pathogens that cause tick-borne diseases (TBDs), including Lyme disease (LD). Several hotspots of elevated LD risk have emerged across Canada as *I. scapularis* expands its range. Focusing on a hotspot in southeastern Ontario, we used high-throughput sequencing on whole ticks and dissected salivary glands and midguts. Compared to whole ticks, analysis of salivary glands and midguts revealed greater β-diversity among microbiomes that are less dominated by *Rickettsia* endosymbiont bacteria and enriched for pathogenic bacteria including a *Bbsl-* associated *Borrelia*, *Borrelia miyamotoi*, and *Anaplasma phagocytophilum*. We also find evidence of co-infection of *I. scapularis* in this region by multiple microbes. Overall, our study highlights the challenges and opportunities associated with the surveillance of the microbiome of *I. scapularis* for pathogen detection using metabarcoding and metatranscriptome approaches.

## Introduction

One-fifth of emerging human infectious diseases are transmitted by arthropod vectors, forty percent of which are transmitted by Ixodidae ticks, representing an increasingly large threat to human health and welfare (1, 2). Ticks in the genus *Ixodes* are important vectors of human disease, including LD, tick-borne relapsing fever, anaplasmosis, babesiosis, ehrlichiosis, tick-borne encephalitis, and Powassan virus (3–6). In Eastern North America, *I. scapularis* is the primary vector of LD, contributing to the more than 300,000 estimated cases in the United States of America (USA) each year (7, 8). In addition to *B. burgdorferi* sensu stricto (s.s.), *I*. *scapularis* is also a vector for the broader *Bbsl* complex of spirochete pathogens, including *B. mayonii*, *B. kurtenbachii*, *B. bissettiae*, and *B. andersonii* (9–14).

A variety of co-occurring pathogens have been detected in *I. scapularis*, including *Bbsl* spirochetes, as well as *A. phagocytophilum, B. miyamotoi, Babesia microti, Ehrlichia muris eauclarensis*, and Powassan virus (15). The large number of pathogens harbored by *I. scapularis* increases the potential for polymicrobial infections, which can complicate diagnosis and result in greater severity of disease (16). While advances in high-throughput sequencing (HTS) have enabled broader detection and monitoring of co-occurring pathogens and microbes harbored by ticks, this technique is not often used for tick monitoring in Canada.

Experimental dysbiosis of the tick microbiome has revealed complex interactions among the bacterial community that can influence pathogen colonization in the midgut (17–20). It has also been shown that *I. scapularis* produces antimicrobial factors that can modulate microbe colonization (20–22). Given the complex interactions that occur in the midgut and the salivary glands of *I. scapularis*, further screening of these tissues from field-collected ticks using HTS can provide better resolution of the interplay between pathogen and tick microbiome at a tissue-specific level.

In addition to immunological factors, the effects of ecological interactions within the tick microbiome remain poorly understood in *I. scapularis*. Recent studies in *I. scapularis* have revealed maternally-transmitted endosymbiotic bacteria, *Rickettsia buchneri*, in relatively high abundances in female *I. scapularis* compared to males, and at higher levels following engorgement (23–26). A negative correlation in abundances of *B*. *burgdorferi* and *R. buchneri* was found among male *I. scapularis* (27), while another study found a potentially negative correlation between *B. burgdorferi* and ‘environmental bacteria’ including *Bacillus* sp., *Pseudomonas* sp., or uncharacterized Enterobacteriaceae (28), implying potential for competition or some other form of interaction to occur between different bacterial associates. Experimental work has shown that *R. buchneri* inhibits the growth of other pathogenic Rickettsiaceae in tick cell lines (29), which may be related to the expression of cryptic genes for interbacterial antagonism (30). Such findings suggest that endosymbionts may influence TBD risk, but to our knowledge correlations between *R. buchneri* and microbial pathogens have not been assessed in wild populations of *I. scapularis* in Canada.

Although most are not known as human pathogens, there are reported cases of tick endosymbiont transmission to humans (31–34). Indeed, many tick endosymbionts are closely related to human pathogens, with examples of evolutionary transitions between pathogenic and symbiotic lifestyles occurring within the Spotted Fever Group of *Rickettsia* (35, 36), *Coxiella* spp. (37), and *Francisella* spp. (38). These bi-directional transitions represent a symbiont-pathogen continuum, further emphasizing the value of HTS surveillance of tick microbiome communities to complement more targeted monitoring initiatives.

While bacteria are commonly studied, ticks also harbor archaea, viruses, and other eukaryotic parasites (16, 39, 40), of which influence on TBD risk remains mostly unknown. Since targeted meta-genome sequencing of the 16S rDNA is specific for bacteria, dual rRNA-depleted metatranscriptome sequencing offers a complementary approach to identify bacterial and non-bacterial associates, as well as capture functional genes expressed within the microbiome. Metatranscriptome studies in *I. scapularis* have revealed cross-kingdom interactions that occur within the microbiome of the tick and may affect TBD risk. For example, non-random co-occurrences have been detected between *Babesia microti* and blacklegged tick phlebovirus (BTPV), *B. burgdorferi* and BTPV (41), and South Bay Virus (SBV) and *B. burgdorferi* (42). More generally, metatranscriptome investigations of tick microbial gene expression have revealed unprecedented diversity of RNA viruses (41–43), including novel pathogens (44, 45).

Recent geographic range expansion of *I. scapularis* is occurring across the USA (46) and into Canada (47–50). Northward expansion of *I. scapularis* ticks into southern Canada is patchy, resulting in regional LD ‘hotspots’ and geographic variation in microbial communities (51–54). Microbiome and transcriptome data from *Ixodes* ticks are rare in Canada but could provide insight into how range expansion shapes microbial communities and the epidemiology of TBDs as part of a transdisciplinary approach advocated by Talbot *et al*. (55). Specifically, metabarcoding and metatranscriptome data from ticks collected in field surveys provide ‘Ecological Evidence’ and ‘Molecular Evidence’ to inform epidemiological models and targets for diagnostic tests. The goal of our study was to understand how tick biology and environment contribute to microbiome composition and pathogen prevalence in natural tick populations, using complementary approaches. Here we present results of HTS-based surveillance of *I. scapularis* microbiomes in a LD hotspot in Canada, using both targeted sequencing of the V4 region of the 16S rDNA gene, and ribosomal RNA-depleted, whole shotgun RNA metatranscriptome sequencing. We examined tissue-specific differences between the salivary gland, midgut, and other internal viscera, as well as whole-tick extractions from three distinct sites representing different land-use histories. Our study demonstrates how HTS methods can help quantify changing health risks associated with range expansion by an arthropod vector of increasing public health concern.

## Results

### Bacterial Community inferred from V4 by 16S rDNA sequencing

The core bacterial community associated with *I. scapularis* was represented by 25 dominant amplicon sequencing variants (ASVs), each with a minimum total relative abundance of greater than 0.1 % (Figure 1; Supplemental Table S1). Within the core, only *A. phagocytophilum* (ASV8) and *B. miyamotoi* (ASV2) were identified at the species level by the *DADA2* pipeline. We identified three primary ASVs of Concern (AoCs) from the core microbiome, which included *Borrelia* sp. (ASV3), *B. miyamotoi*, and *A. phagocytophilum*. Among these AoCs, the potentially LD-causing spirochete, *Borrelia* sp. was the most widely detected across all tissue types and sampling sites in this study, ranging from trace levels up to 98 % relative abundance. We also detected the relapsing fever spirochete, *B. miyamotoi*, at >99 % relative abundance from the salivary gland and midgut of a single tick collected at the Lemoine Point (LP) site.

**Figure 1:**
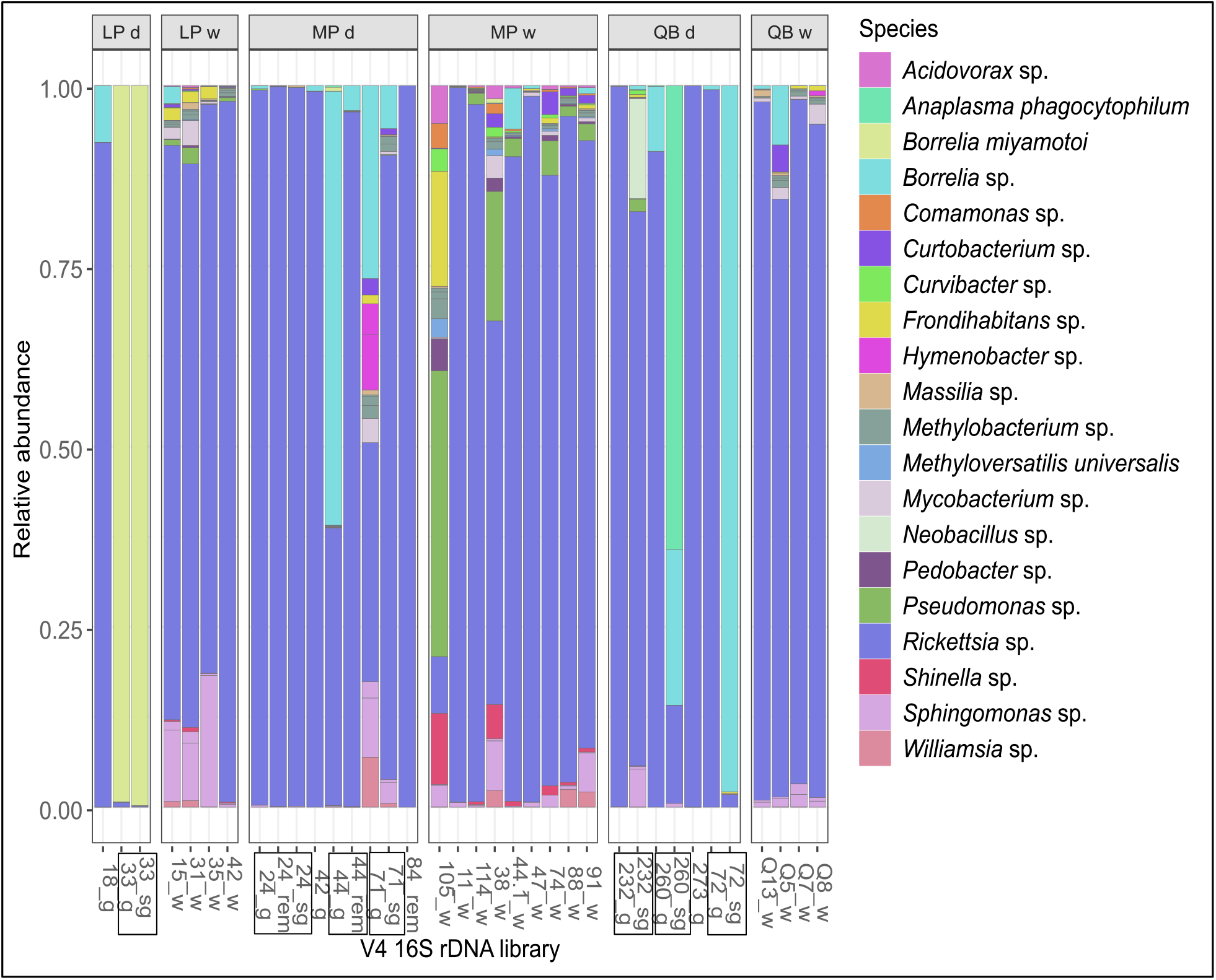
Relative abundance of amplicon sequence variants (ASVs) of the V4 16S rDNA region detected in the core microbiome of *I*. *scapularis*. Core bacteria are defined as ASVs with a minimum relative abundance of 0.1 % in at least one sequencing library. Libraries originating from different tissues of the same individual are denoted by black rectangles around the sample identifiers on the x-axis. Sample types are salivary gland (sg), gut (g), remaining internal viscera (rem), whole (w), or dissected (d) in the library identifiers/ facet titles. Samples were collected from three locations in eastern Ontario: Lemoine Point (LP), Murphy’s Point (MP), and Queen’s University Biological Station (QB) (see facet titles).

Aside from the three AoCs detected in the core microbiome, another major feature of the *I. scapularis* bacterial community was the widespread detection of *Rickettsia* sp. (ASV1), which was common across all three sites from whole-tick and dissected tissue samples. A total of six ASVs were identified as *Rickettsia* sp. (Supplemental Table S2) with the sequences ranging in length from 254 – 258 bp and sharing 93.8 – 99.6 % sequence similarity based on multiple sequence alignment (CLUSTAL O (1.2.4); data not shown). Of these, ASV1 contributed >99.95 % of all *Rickettsia*-associated sequences captured in this study, which shares 100 % homology to the V4 region of the 16S rDNA of *R. buchneri* (endosymbiont of *I. scapularis*). Also notable, *Williamsia* sp. (ASV16) and *Mycobacterium* sp. (ASV11), *Sphingomonas* sp. (ASV4), and *Pseudomonas* sp. (ASV5) were detected in the core microbiome at two or more of the sampling sites (Figure 1).

We consider the detection of ASVs from the sequencing libraries to be reliable. The negative control PCR and bead libraries contained markedly fewer sequences following our data processing but removed one salivary gland-based sample (42_sg) (Supplemental Figure S1), noting that an initial inspection of ASVs detected among the three separate batch PCR replicate libraries determined the presence of several additional *Rickettsia-*associated ASVs almost exclusively from a single PCR batch, including the negative control PCR (Supplemental Table S3 and Supplemental Figure S2). Since these were unexpected results, the entire batch replicate was removed from further analysis, while the two more consistent batch replicates were retained (noting seven of the dissected samples only had a single PCR batch replicate).

From these sequences, 305 potentially contaminating ASVs, including taxa typically known from environmental samples (e.g., *Sediminibacterium, Pseudomonas, Paraburkholderia*, *Enhydrobacter, Massilia*) or human sources (e.g., *Staphylococcus, Streptococcus, Acinetobacter*, *Cutibacterium, Chryseobacterium*, *Corynebacterium*) were identified using *decontam* and removed from the dataset, (Supplemental Table S4). Based on the asymptotes of rarefaction curves, bacterial communities associated with whole ticks generally contained more ASVs than those detected from the various dissected tissues (Supplemental Figure S3). Overall, each library contained 1 to 139 ASVs, with an average of 54.

Sequences attributed to ASV7 were detected in 2 out of 28 of the tick samples, both from salivary gland samples from the Queen’s University Biological Station (QB) site and could not be assigned beyond Kingdom *Bacteria* in our pipeline (Supplemental Table S5). However, a BLAST search of GenBank’s nucleotide database identified similarity (e-value = 3e^-106^; 248/264 identities) to the apicoplast of *Babesia* sp. Dunhaung (MH992225.1) and *Babesia* sp. Xinjiang complete genome (MH992224.1) and Kashi isolate (KX881914.1) (56). Phylogenetic analysis placed ASV7 within the Clade X *Babesia* “sensu stricto” (i.e., classical or true *Babesia*) (Supplemental Figure S4). The *Babesia*-associated ASV7 was not included in further analysis.

Co-infection of ticks with multiple AoCs were identified in the salivary gland of a tick from the QB site (260_sg). This sample was coinfected with *Borrelia* sp. and *A. phagocytophilum*, representing 22 % and 64 % relative abundance, respectively (Figure 1). In contrast, the midgut of the same individual (260_g) contained 9 % relative abundance of *Borrelia* sp. and trace levels (< 1 % relative abundance) of *A. phagocytophilum*. Moreover, the *Babesia* sp. classified as ASV7 (see above) was detected among other bacterial AoCs, either co-infecting the salivary gland with *Borrelia* sp. in sample 72_sg, or present in a triple infection with both *Borrelia* sp., and *A. phagocytophilum* in 260_sg, both from salivary gland collected at the QB site.

Additional phylogenetic analysis of the 258 bp V4 16S rDNA sequence from *A. phagocytophilum* was 100 % identical to sequences from strain Ap-var-1, which was previously detected in Canada, and distinct from the known pathogenic strain Ap-ha also detected in Canada (57) (Accession: HG916767.1; Supplemental Figure S5). The sequence from *Borrelia* sp. matched 100 % with the partial 16S rRNA sequence from *B. bissettiae* DN127 (255/255 bp identities; NR_114707.1) (58) and *B. burgdorferi* s.s. isolate 15-0797 (255 / 255 bp identities; MH781147.1) (Supplemental Figures S6). The V4 16S rDNA sequence from *B. miyamotoi* matched 100 % with the *B. miyamotoi* strain HT31 (NR_025861) with 257 / 257 bp identities, which is distinct from the *Bbsl* complex (Supplemental Figures S6).

We used LEfSe to explore variation in ASV abundance among tissue types and sample sites. The abundance of two AoCs of concern, *Borrelia* sp. and *A. phagocytophilum* discriminated between dissected tick tissues and wholes-tick samples, based on LDA scores (Supplemental Figure S7A). The third AoC, *B. miyamotoi,* discriminated between tissue samples from the LP site only (Supplemental Figure S7B). While only the abundance of the three AoCs discriminated among dissected tick tissues, the LEfSe identified several core ASVs that discriminate among sites for the whole-tick samples (Supplemental Figure S7B). Specifically, *Methylobacterium* sp., *Mycobacterium* sp., *Curtobacterium* sp., *Massilia* sp., *Hymenobacter* sp., and Order *Rhizobiales* were more common at the QB site; *Pseudomonas* sp., and *Shinella* were more common at the MP site; and *Sphingomonas* sp., *Frondihabitans* sp., and *Clavibacter* sp. were more common at the LP site.

Consistent with the LEfSe, Principle Co-ordinates Analysis (PCoA) of the V4 16S rDNA libraries using weighted UniFrac distances separated samples primarily by high abundance of AoCs, which explain 89.5 % of the total variation among the bacterial communities associated with *I. scapularis* (Figure 2). Samples defined by a high relative abundance of *Rickettsia* sp. clustered together (bottom-left corner in Figure 2) whereas those with relatively high abundances of *Borrelia* sp., *B. miyamotoi*, *A. phagocytophilum*, or *Pseudomonas* sp. were separated from the main cluster of samples.

**Figure 2:**
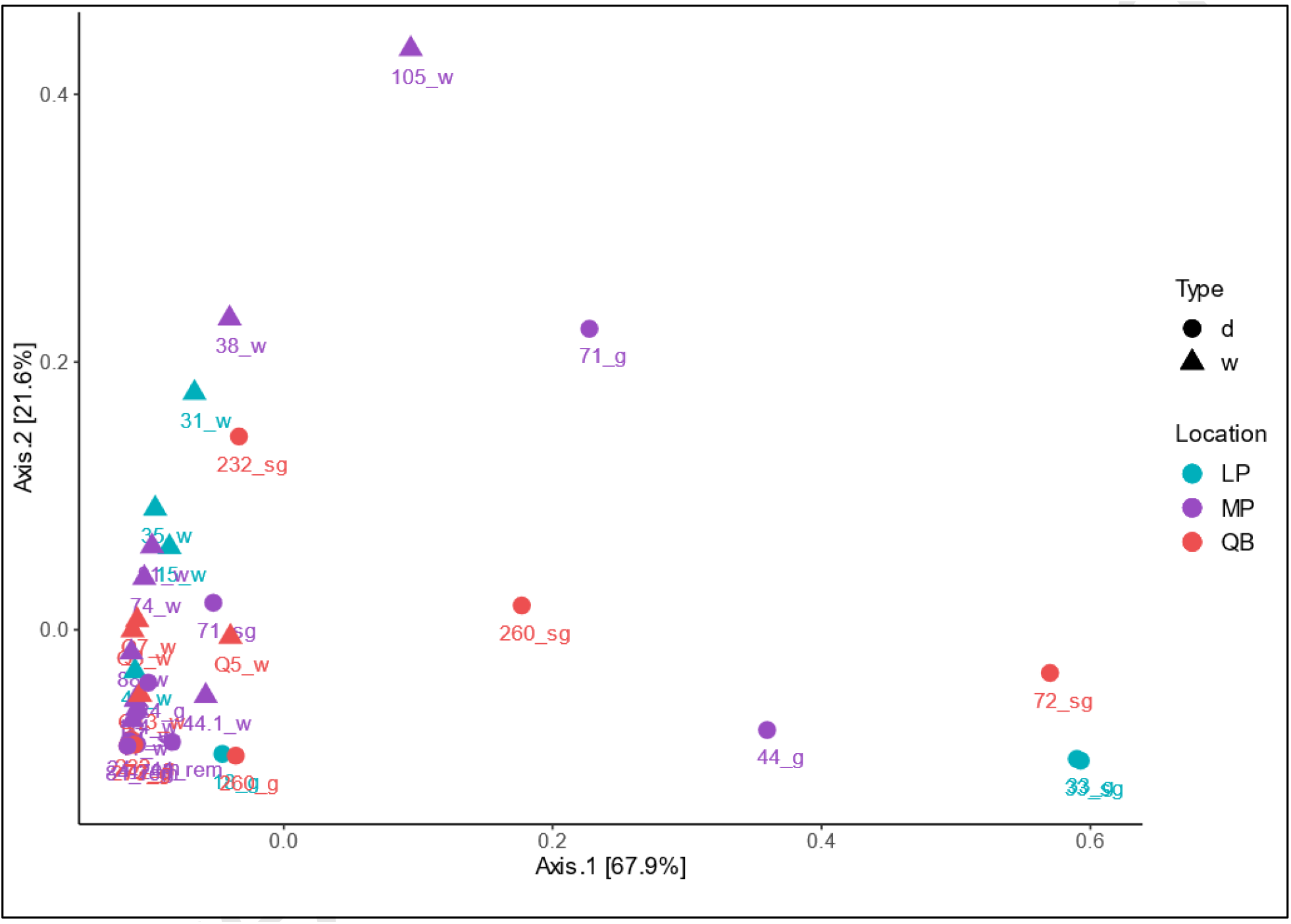
Principle Co-ordinates Analysis (PCoA) of weighted UniFrac distances for ASVs detected from *I. scapularis* using V4 16S rDNA sequencing (all ASVs were agglomerated at the lowest taxonomic level; N = 419). Sample types are indicated as salivary gland (sg), gut (g), internal viscera (rem), or whole (w) in library names. Samples were collected at three locations, Lemoine’s Point (LP), Murphy’s Point (MP), and Queen’s University Biological Station (QB). Shapes indicate tissue type - whole (w) or dissected (d).

In contrast to weighted UniFrac, the first two axes of the unweighted UniFrac distances explain only 34.1 % of the variation in the data (Figure 3A). Generally, the unweighted UniFrac ordination separated the bacterial communities by tissue type, with whole-tick samples clustering away from dissected samples, with the exception of a few samples (71_g, 71_sg, Q5_w, Q13_W, and 47_w) found in ellipse peripheries where the ellipses between whole and dissected types overlapped. Overall, we detected 409 ASVs from the ticks and tissues that were sampled, with 112 ASVs unique to whole ticks, 152 unique to dissected tissues, and 145 shared between them (Figure 3B). Among these uniquely identified ASVs, strains of bacteria in the Phylum Verrucomicrobia were more common in whole-body samples from the LP and MP sites, compared to strains in the Phylum Firmicutes, which were more common among dissected tissues collected across all three sampling locations (Supplemental Figure S8).

**Figure 3:**
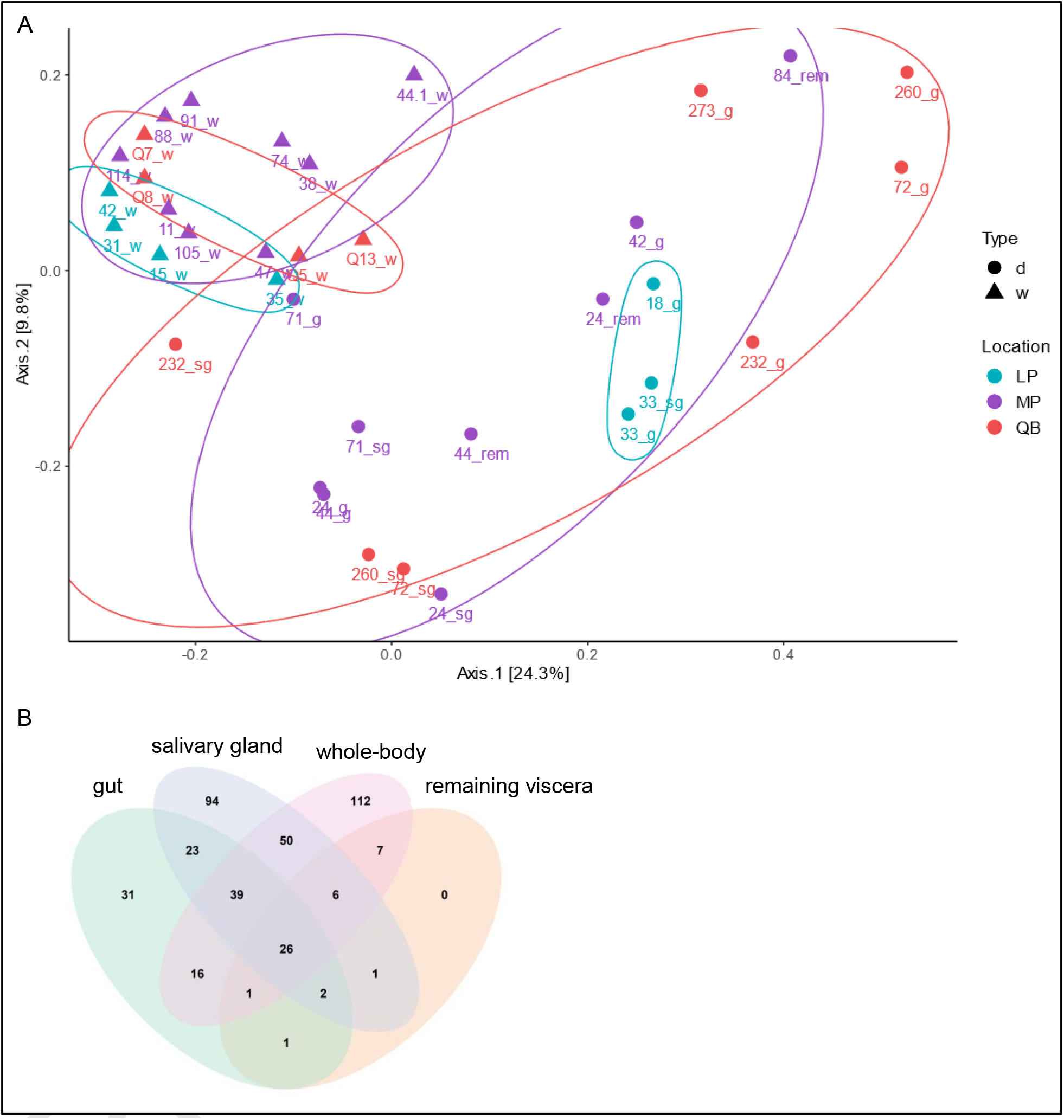
A) Principle Co-ordinates Analysis (PCoA) of unweighted UniFrac distances for ASVs. The tissue types are indicated in the library names as either dissected (d): salivary gland (sg), gut (g) and internal viscera (rem), or whole (w). Location abbreviations: Lemoine’s Point (LP), Murphy’s Point (MP), and Queen’s University Biological Station (QB). B) Venn diagram of ASVs detected using V4 16S rDNA sequencing from the whole *I. scapularis* or dissected salivary gland, midgut, and other remaining internal viscera.

Differences in β-diversity between communities within individual female ticks sampled from whole or dissected tissues were identified. The average unweighted UniFrac distance of 0.512 (SD = 0.078) between whole tick libraries was significantly lower (*P* < 2.22e^-16^) compared to the average distance of 0.707 (SD = 0.118) among libraries from dissected ticks (Supplemental Figure S9). Furthermore, the average distance among the whole tick libraries was also significantly lower (*P* < 2.22e^-16^) than the average distance of 0.706 (SD = 0.104) measured from libraries between the groups. Using a permutation test for homogeneity of multivariate dispersions, we also identified a significant difference in variance between whole and dissected sample types. The unweighted UniFrac PCoA was repeated using only the top 500 most relatively abundant ASVs, which was consistent with the previous findings that used ASVs agglomerated at the lowest taxonomic level (Supplemental Figure S10A) but the PCoA inverted Axis 2 and showed slightly more distinction between dissected and whole tick samples (Supplemental Figure S10B).

### Whole-shotgun RNA sequencing

Following the removal of transcripts associated with unwanted phyla (see methods), we further analyzed the remaining 402 transcripts of which, endosymbiont *R. buchneri* comprised over 75 % (Table 1). Single-stranded negative-sense RNA viruses were common in the *I. scapularis* metatranscriptome (Figure 4), including transcripts for N and L proteins from two different viruses in the family *Bunyaviridae*. Of these, a South Bay Virus (SBV) from the genus *Nairovirus* was found to be highly expressed in all four tick samples. We also identified transcripts from two variants of blacklegged tick phlebovirus (BLTPV), from the genus *Phlebovirus* of *Bunyaviridae*, and from three different variants of the currently unclassified *Ixodes scapularis*-associated virus (ISAV) (Figure 4). Finally, a partial N protein transcript of 508 bp length shared sequence similarity with the Chimay rhabdovirus N protein by BLASTx (e-value = 1.69e^-17^), but it was expressed at low levels relative to the other virus-associated transcripts.

**Figure 4:**
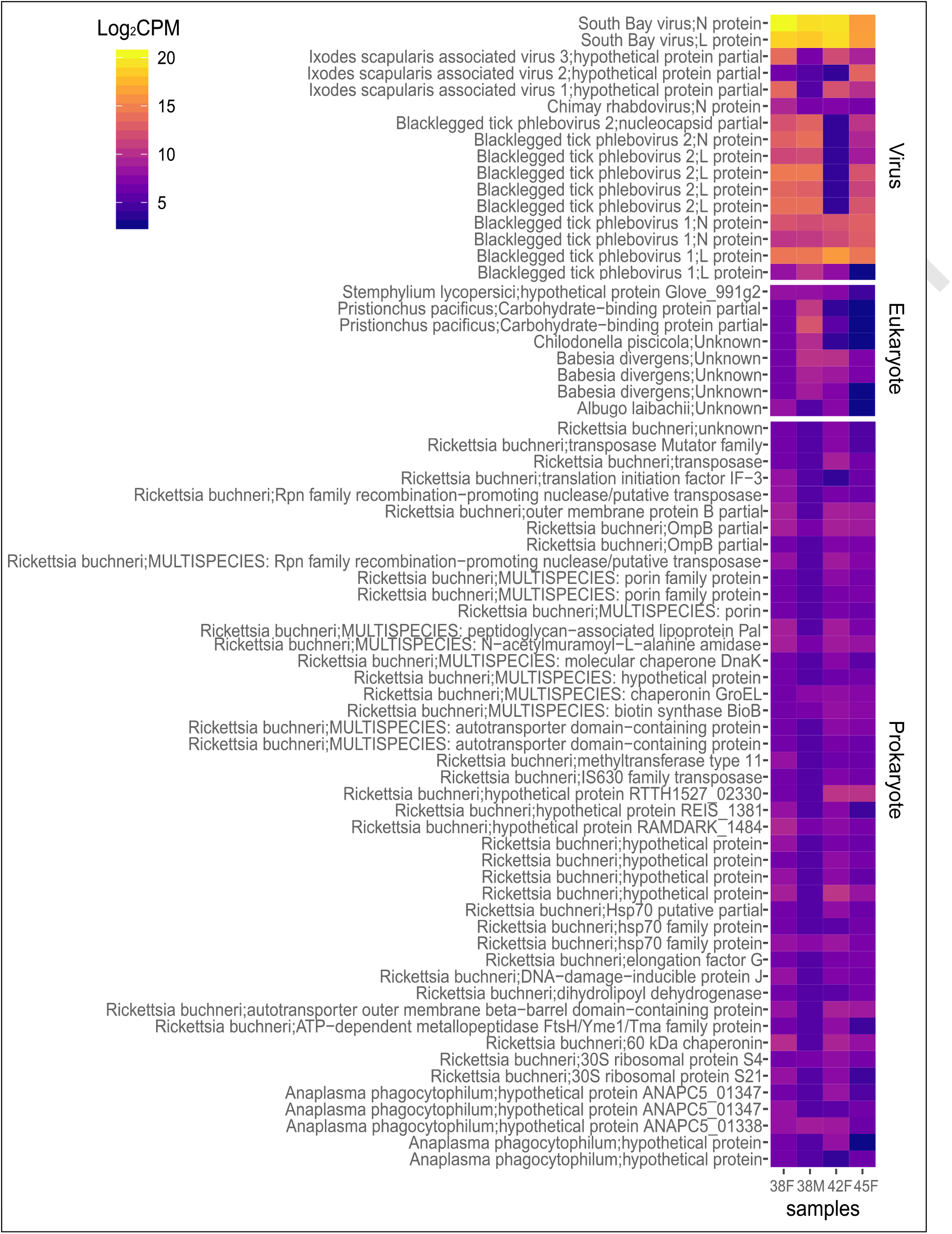
Log_2_ counts-per-million (CPM) expression of non-*Ixodes scapularis* assembled transcripts from transcriptome libraries of female (38F, 42F and 45F) and male (38M) adult *I. scapularis*.

**Table 1:** Proportion of taxonomic assignments in the I. scapularis metatranscriptome

| Super kingdom | Annotation | Proportion (%) |
| --- | --- | --- |
| Eukaryota | Other non-arthropod | 10.70 |
| Prokaryota | *R. buchneri* | 75.12 |
| Prokaryota | *R. parkeri* | 0.50 |
| Prokaryota | *B. burgdorferi* | 1.24 |
| Prokaryota | *B. miyamotoi* | 0.25 |
| Prokaryota | *A. phagocytophilum* | 3.98 |
| Prokaryota | Other | 3.48 |
| Viruses | Viruses | 4.73 |

Transcripts of potential eukaryotic origin, including some with the highest BLAST hits against the tick-vectored intracellular parasite *Babesia divergens*, were identified in low abundance relative to the virus-associated sequences. Transcripts of prokaryotic origin were identified at intermediate abundance, including *R. buchneri* and *A. phagocytophilum*, while *Borrelia*-associated transcripts were captured at relatively lower abundances. Forty *R. buchneri* genes were expressed in *I. scapularis*, including genes encoding for *bioB* (involved in biotin synthesis), outer membrane proteins, lipoproteins, peptidoglycan modification, autotransporters, porins, heat-shock proteins, transposases, molecular chaperones, DNA methylation, ribosome-associating proteins and other hypothetical genes of unknown function.

## Discussion

Using a multi-omics approach with HTS, we found evidence for endosymbiotic and pathogenic microorganisms of viral, eukaryotic, and prokaryotic origin associated with adult *I. scapularis* collected from a LD hotspot located in southeastern Ontario. Consistent with other 16S rDNA-based sequencing studies of *I. scapularis*, the most common and relatively abundant bacteria found in our study belong to the genus *Rickettsia*. In addition, we detected three primary AoCs, including *A. phagocytophilum*, a spirochete from the *Bbsl* complex (*Borrelia* sp.), and non-*Bbsl* relapsing fever spirochete *B. miyamotoi*. Using whole-shotgun RNA sequencing we found that *I. scapularis* was infected with the three primary AoCs in the preceding sentence, as well as *Babesia* sp., and *R. buchneri*, though few pathogen-associated transcripts were captured from four adult ticks that were sequenced. A diversity of RNA viruses, especially from the family *Bunyavirales*, were identified in relatively high abundances in the metatranscriptome. Our findings highlight the usefulness of combining both non-targeted meta-transcriptomics and tick organ-specific 16S rDNA sequencing to assist in the detection of pathogens and polymicrobial infections involving diverse pathogenic and symbiotic microbes of the *I. scapularis* microbiome.

### Potential pathogens

In contrast to targeted surveillance, metabarcoding by HTS is a cost-effective method to simultaneously detect diverse *Bbsl* and non-*Bbsl Borrelia* in ticks, which can inform risk assessment of TBDs in Canada. Using HTS, our study revealed *Borrelia* sp. from the *Bbsl* complex at three sampling sites, and at higher abundance in the dissected midgut and salivary gland tissues compared to whole tick samples. We also detected the non-*Bbsl* relapsing-fever spirochete, *B. miyamotoi* in relatively high abundance from the dissected tissues of a single tick and in other sites at a lower frequency than *Bbsl*-associated ASV. Although *B. burgdorferi* s.s. was first recognized as the genospecies responsible for LD, several other pathogenic *Bbsl* genospecies that cause Lyme, or other Lyme-like diseases are also transmitted by *I. scapularis*, including *B. mayonii*, *B. kurtenbachii*, *B. bissettiae,* and *B. andersonii* (9–14). Beyond the *Bbsl* complex, the causative agent of relapsing fever, *B. miyamotoi*, has been found in *Ixodes* species from other parts of North America (59).

Few studies from Canada have used HTS, but Sperling *et al.* (60) found that *Borrelia*-positive ticks had a higher α-diversity compared to *Borrelia*-negative ticks, and a high variance in *Borrelia* sequence abundance (0 – 93 %) from ticks identified as *Borrelia*-positive using qPCR (60). In another study of the bacterial communities of adult *I. scapularis* from eastern and southern Ontario, Canada, Clow *et al.* (61) identified *Borrelia* from 70 % of the ticks tested with an average relative abundance of 0.01 %. Similarly, we also identified a low abundance of *Borrelia* spp. from whole ticks, but the same sequences, when detected from some individual tick midgut and salivary glands were found in much higher relative abundance compared to whole ticks. Our analysis highlights the value of metabarcoding the V4 16S rDNA from tick salivary glands and midguts to identify potential pathogens, and for discrimination between *Bbsl* and non-*Bbsl Borrelia*, which has implications for modelling risk associated of TBDs in Canada.

We found another potentially important human pathogen, *A. phagocytophilum*, at low frequency in this study, mainly associated with ticks collected from the QB site. Such findings are consistent with other PCR-based screening that detected *A. phagocytophilum* infection frequencies from 0.4 to 8.9 % in *I. scapularis* from Canada (62–64). Our work also demonstrates the value of metabarcoding by HTS in a phylogenetically-informed analysis to resolve the strain-specific distribution of this potential pathogen. Specifically, we find strain Ap-var-1, which differs from the known pathogen strain Ap-ha (57), and although the Ap-var-1 strain is not known to be pathogenic, human health risk assessments may be better informed through greater sampling effort across more sites in the region to capture potentially virulent strains that may be present at lower frequencies within *I. scapularis* populations.

In addition to distinguishing between three bacterial AoCs, we find that V4 16S rDNA sequencing also contained *Babesia*-associated apicoplast sequences from the dissected salivary gland of two ticks collected from the QB site. Interestingly, the *Babesia-*associated sequence was co-associated with either *Borrelia* sp., or *Borrelia sp*. and *A. phagocytophilum*. While this *Babesia* sp. is not closely related to human pathogenic *Babesia microti* based on BLAST hits, its pathogenic potential is unknown. It is reasonable to suspect *Babesia odocoilei* because it has been detected in eastern Canada before (65–70), but we are not able to confirm this since the apicoplast sequence is not available for *B. odocoilei*. A next step may include using targeted PCR, metabarcoding with *Eukaryote*-specific primers, or whole metagenomics to further classify this co-associated *Babesia* sp. detected from the QB site.

### Tissue-specific bacterial associates

Our findings of two *Borrelia* species, *A. phagocytophilum*, and *Rickettsia* sp., are consistent with other studies that have also examined bacterial communities associated with specific tissues dissected from *I. scapularis*. *Rickettsia buchneri* is the most common and abundant bacterial species in all organs of female ticks, which is consistent with two other HTS studies (28, 71). Furthermore, Al-Khafaji et al. (72) recently found clear evidence of rickettsial infection in *I. scapularis* SG samples by confocal microscopy. Beyond *R. buchneri*, ours and one other study found *Anaplasma* sp. and two different *Borrelia* spp. to be relatively abundant in the salivary glands and midgut of adult female *I. scapularis* (71). Ross et al. (28) using LEfSe analysis showed that bacteria from the orders *Spirochaetales* and *Rickettsiales* were most likely to explain differences in the bacterial communities associated with tick internal viscera, compared to external washes, which is similar to our LEfSe results, in which *A. phagocytophilum* (Order *Rickettsiales*), and two different *Borrelia*-associated ASVs (Order *Spirochaetales*) discriminated between the communities associated with salivary glands and midgut tissues, compared whole tick samples. Taken together, these studies and ours show that ASV abundance from whole-body tick samples tends to under-represent the prevalence of potential pathogens relative to dissected gut and salivary glands. Although labor-intensive, micro-meter dissections increase sensitivity of pathogen detection, which could help to better model risks posed by TBDs transmitted by adult *I. scapularis*.

HTS sequencing of tick tissues poses two additional challenges. First, the smaller amount of extracted DNA from tissue-specific sequencing is more prone to contamination (28). Second, the β-diversity measures of low biomass samples are more prone to inflation due to uneven sampling depth, whereas samples with relatively fewer sequences have greater uncertainty associated with rare ASVs (73–75). To address these issues, we applied a two-step approach to screen contaminating ASVs from the sequencing data, followed by a rarefaction analysis to assess for sufficient capture of bacterial diversity (supplemental Figure S4). Even after these corrective measures, we identified a greater number of unique ASVs associated with bacterial communities from dissected tissue compared to whole-body samples. This does not appear to be an artifact of small sample size or uncertainty associated with rare ASVs because α-diversity in our lower biomass samples (i.e., dissected tissues) was not inflated relative to the whole-body samples in our rarefaction analysis (supplemental Figure S4). The biological significance of higher diversity in tick tissue is worthy of future study.

### Other microbial associates of interest

Although several pathogens were identified in our study, the most abundant ASVs were *Rickettsia* sp., consistent with previous studies (28, 60, 61, 71, 76–78). Additionally, we found a high relative abundance of *Rickettsia* sp. in the midgut and several salivary gland tissues of several ticks (Figure 1). In *I. scapularis*, *R. buchneri* can colonize the midgut and salivary glands where it secretes proteins into the tick’s saliva (72). Our meta-transcriptomic analysis confirms active transcription of at least one *R. buchneri* biotin synthesis gene, *bioB,* which is encoded on a plasmid along with all the genes for biotin synthesis (79). Our analysis also identified several *R. buchneri*-associated transcripts coding for known secreted factors previously identified by Al-Khafaji *et al.*(72), including transcripts for a peptidoglycan-associated lipoprotein, and moonlighting proteins DnaK and GroL. Beyond the expression of genes for these known secreted proteins, we also identified *R. buchneri*-associated transcripts coding for various secretion systems, including porins, autotransporters, and autotransporter domain-containing proteins. Although *R. buchneri* is not a known human pathogen, our results imply that *R. buchneri* could play a role in tick feeding in ways that may interact with the immune system of the vertebrate host via transmission of rickettsial-secreted factors in tick saliva and gut contents.

Consistent with surveys of *I. scapularis* in other geographic regions (42, 80, 81), we detected *Bunyavirales*-like viruses such as SBV, BLTPV-1, and BLTPV-2, as well as three unclassified ISAVs from the metatranscriptome of *I. scapularis*. These are tick-specific viruses that are not known to be pathogenic to humans but could influence tick physiology and behavior. In contrast to the metatranscriptome reported by Cross *et al.* (42), in which they rarely observed BLTPV-1 and BTPV-2 co-occurring in the same *I. scapularis*, our study detected co-occurrence of both BTPV-1 and BTPV-2 in three out of four of the ticks analyzed, suggesting that both variants can co-exist within the same host.

## Conclusions

Among the female *I. scapularis* that we surveyed using 16S rDNA sequencing, *Bbsl* and non-*Bbsl Borrelia,* and *A. phagocytophilum* were found in relatively high abundances, especially within the salivary glands and midgut. Beyond potentially pathogenic bacteria, 16S rDNA and metatranscriptome analysis also identified *Babesia* sp. of unknown virulence, while metatranscriptome captured a diversity of *Bunyavirales*-like and other unclassified tick-specific viruses, some with relatively high expression. These results demonstrate the added value in combining both 16S rDNA and metatranscriptome sequencing for a more comprehensive view of the entire microbiome of *I. scapularis*, with a dynamic complex of pathogen and endosymbiotic microbes, including bacteria, viruses, and intracellular parasites.

## Methods and Materials

### Tick collection and nucleic acid extraction

Adult *I. scapularis* were collected at the Lemoine Point (LP) Conservation Area (44°13’ 56.228”N; 76° 36’ 45.795”W) within the municipality of Kingston Ontario, Murphy’s Point (MB) Provincial Park (44° 46’ 54.7098”N; 76° 14’ 30.1122”W), and Queen’s University Biological Station (QB) (44° 34’ 4.6524”N; 76° 19’ 57.0966”W). Samples were collected from the QB site in 2016, and from all three of the sites in 2017. Questing adults were captured by flagging with 1 m^2^ white flannel fabric attached to an aluminum bar along a transect of 25 m, checking for attached ticks at 5 m intervals. Ticks were removed by hand or with forceps, placed in 2 mL screwcap tubes containing 70 % ethanol, and later stored at -20 or -80 °C. Prior to extraction, ticks were submerged in a 1 % bleach solution for 30 s and rinsed with lab-grade water. Different extraction methods were used for the DNA of internal tissue, DNA of whole ticks, and RNA of whole ticks, as follows.

We dissected twelve female ticks to remove internal tissue from the exoskeleton and to separate the salivary glands and midgut from the remaining internal viscera for separate extraction and sequencing. The midgut and salivary glands were successfully isolated in ten ticks. Internal tissues were separately macerated in 50 μL of extraction buffer containing 1 mM EDTA, 25 mM NaCl, 10 mM Tris-HCl pH 8.0, and 200 μg mL^-1^ Proteinase K (VWR). The DNA was purified using solid phase reversible immobilization (SPRI) beads at 2.5 ✕ volume and resuspended in 20 μL laboratory-grade water.

In addition to the dissected ticks, we extracted DNA from seventeen whole female ticks. Following sterilization as above, these ticks were dried at room temperature, frozen to -80 °C and pulverized in a Next Advance Bullet Blender Storm 24 at 100 Hz in 2 mL tubes containing equal volumes of 0.2- and 0.5-mm low binding ZrO beads (SPEX®) for 3 - 4 min, re-freezing every 1 min until no large visible fragments remained. Next, each sample was incubated with 500 µl of pre-heated Cetyl trimethylammonium bromide (CTAB) buffer (100 mM Tris-HCl pH 8.0, 25 mM EDTA, 1.5 M NaCl, 3 % CTAB, 1 % polyvinylpyrrolidone, and 1 % β-mercaptoethanol) at 62 °C for 30 min. To isolate the DNA, 500 µl of chloroform was added, the samples were centrifuged at 1,300 x g for 15 min and then the supernatant was transferred to a new microcentrifuge tube. The DNA was precipitated in 1 mL pre-chilled 100 % ethanol, mixed by inversion and then incubated at -20 °C for 30 min. The DNA was pelleted by centrifuge at ∼21,000 x g for 15 min, washed twice with 1 mL 75 % ethanol, and resuspended in 15 µl laboratory-grade water.

We isolated RNA from one male and three female ticks using the same sterilizing and pulverizing steps as the whole-tick DNA protocol above but with 500 µL TRIzol Reagent (Invitrogen) following the manufacturer’s protocol and resuspending purified RNA in 15 μL lab-grade water.

### Library preparation

The V4 region of the 16S rRNA gene was amplified from tick DNA using forward 515F (5’-GTGCCAGCMGCCGCGGTAA-3’) and reverse 806R (5’-GACTACHVGGGTWTCTAAT-3’) primers (82). The PCRs were completed in triplicate using a two-step approach, with each replicate including a control PCR (template replaced with PCR-grade water). The first-step PCRs were completed in 25 µL volumes using ∼ 2 - 3 ng of DNA template, 2 U Platinum^TM^ *Taq* polymerase high-fidelity (Invitrogen), 0.5 µg µL^-1^ bovine serum albumin (NEB), 3 mM MgSO_4_, 200 µM dNTPs, and 0.4 µM each 515F and 806R in 1 x High Fidelity PCR Buffer. Using a SimpliAmp^TM^ thermal cycler (Applied Biosystem), the reactions were incubated at 94 °C for 3 min, followed by 20 cycles of 94 °C for 45 s; 53 °C for 1 min; and 72 °C for 1 min 30s, and then a final elongation at 72 °C for 10 min. These first-step products were purified using Sera-Mag Magnetic SpeedBeads according to manufacturer directions, and 15 µL of this purified PCR product was used as the template for the second-step PCR for 12 cycles as above in 50 µL volumes. The second-step reactions were cycled 12 times using the above program with the 515F/806R phasing primers (83) to add the appropriate Illumina sequencing adapters, barcodes, and heterogeneity spacer components.

Following the second-step PCR, the products were then purified with SPRI beads as described above and the concentration of the resultant DNA was quantified using the dsDNA High Sensitivity Kit (DeNovix) on a DS-11 FX Spectrophotometer / Fluorometer (DeNovix) and pooled for equivalency. Twenty-four libraries failed to amplify and were excluded from the sequencing pool. Paired-end sequencing was performed at Genome Quebec using the MiSeq PE300 (2 x 300 bp) Platform (Illumina, V3 Kit). A Small Read Archive of this sequencing data is available on GenBank under Accession: SRR17194087.

Tick RNA was assessed using the 2100 Bioanalyzer (Agilent) to ensure sufficient quantity and quality before proceeding to rRNA depletion using the NEBNext rRNA Depletion Kit (Human/Mouse/Rat). Next, the sequencing libraries were prepared using the NEBNext Ultra II Directional RNA Library Prep Kit for Illumina with NEBNext Multiplex Oligos for Illumina, and the resulting DNA concentrations were determined with the NEBNext Library Quantification Kit, all according to NEB’s protocols. Finally, the cDNA libraries were sequenced at the Infectious Disease Sequencing Lab in the Kingston Health Sciences Centre using the MiSeq platform with a MiSeq reagent kit v2 (300 cycles) (Illumina MS-102-2002) producing 150 bp paired-end sequencing data. A Sequence Read Archive of this sequencing data is available on GenBank under Accession: SRR17194086.

### 16S rRNA sequencing and analysis

Raw sequence data were demultiplexed using cutadapt (v2.8) allowing up to 10 % maximum error within the barcoded region (84). Libraries were demultiplexed based on barcode and heterogeneity spacer sequences and re-labeled using the multiple move tool *mmv*. The PCR reactions that failed to amplify during library prep were omitted from the data set prior to analysis. The remaining libraries were then processed and analyzed with a variety of packages for R (v4.1.0) (85) in RStudio (v1.2.5033) (86) using a reproducible analysis pipeline available at https://github.com/damselflywingz/tick_microbiome (also see data availability statement below).

Forward and reverse libraries were filtered with default settings and trimmed using *DADA2* for R (v1.20.0) (87). The unpaired forward and reverse libraries were each trimmed to a final length of 220 bp, after 5’ primer removal and trimming for poor quality base-calls in the 3’-regions. Next, a core sample inference denoising algorithm was applied to the pooled libraries (87). Following denoising, the forward and reverse pairs were merged and filtered for lengths between 250 – 265 bp (V4 region using 515F/806R is 254 bp). Identical sequences were collapsed into the longest representative sequence and screened for chimeras using the consensus method of *de novo* bimera removal.

Amplicon sequence variants (ASVs) were taxonomically assigned by implementing Ribosomal Database Project (RDP) (88) naïve Bayesian classifier (89) using the *DADA2*-formatted training set 18 from version 11.5 of the RDP (downloaded August 26, 2021). Species-level assignments were determined by comparing ASVs against the *DADA2*-formatted RDP species-assignment training set to identify unambiguous exact matches (90). The 16S V4 rDNA dataset was converted into a phyloseq object and analyzed with *phyloseq* (v1.36.0) (91, 92).

Contaminant ASVs were removed with *decontam* (v1.12.0) by first applying the prevalence method on individual PCR batch replicate libraries and then applying the frequency method on the whole dataset (93). During preliminary analysis, the libraries associated with one out of the three PCR batch replicates were found to contain 31 additional *Rickettsia-*associated ASVs that were not identified from the other two replicates. This problematic replicate was considered atypical compared to the other two PCR batch replicates and was subsequently removed from the analysis. Inconsistencies between low abundance ASVs across PCR replicate libraries were also removed by retaining only ASVs occurring in the remaining two replicate libraries unless only one library was available (n = 7). Libraries that were generated from any remaining replicate PCRs were combined *in silico*.

Any ASVs assigned at the Phylum level within Kingdom Bacteria were retained, except any ASVs assigned as ‘‘*Cyanobacteria*/*Chloroplast*’ at the Phylum level, which were removed. Only ASVs with counts greater than four were retained for further analysis. Any sample libraries with 7,500 or fewer ASVs were removed. Rarefaction curves were generated using *ranacapa* (v0.1.0) (94). The core bacterial community of *I. scapularis* was defined as the set of ASVs with ≥ 0.1 % relative abundance in at least one library. Relative and total abundances of the core bacterial community were visualized with *ggplot2* (v3.2.1) (95).

Phylogenetic reconstructions of potential pathogenic *Borrelia*- and *A. phagocytophilum-* associated ASVs identified from the tick core microbiome were generated using 16S rDNA sequences available from NCBI 16S rRNA reference database (accession: PRJNA3175; downloaded November 16, 2021) (96), and select sequences available through NCBI’s GenBank website (97). The *Phangorn* R package (v2.7.1) was used to generate optimized phylogenies for these ASVs (98). For each phylogeny, a maximum-likelihood tree was generated using a pre-optimized distance-based method (i.e., rooted or unrooted) and the best substitution model was identified with *Phangorn*. The branch confidence of each of the final phylogenetic reconstructions was tested with 1,000 bootstraps.

To assist with the taxonomic identification of known pathogens, we obtained 27 sequences of *Borrelia* and *Borreliella* 16S rDNA-associated sequences from the NCBI 16S rRNA reference database (96), and an additional sequence from *B. burgdorferi* isolate 15−0797 (Accession: MH781147.1) that did not contain any ambiguous. Potential *Borrelia* sequences from the above pipeline were aligned to these reference sequences using a staggered alignment with *DECIPHER* in R (v2.20.0) (99). Similarly, we used eleven *Anaplasma* and *Ehrlichia* sequences from GenBank’s 16S rRNA reference database and two additional V4 16S rDNA sequences from strains of *A. phagocytophilum* that were previously detected in Ontario (57). Of these, the Ap-ha (Accession: HG916766.1) strain is considered pathogenic, while Ap-var 1 (Accession: HG916767.1) is not known to be pathogenic (57).

Next, ASVs associated with the entire bacterial community were agglomerated at the lowest level of taxonomic identification. Linear discrimination analysis (LDA) Effect Size (LEfSe) comparing 16S V4 rDNA libraries from 1) whole versus dissected tissue types across all sites, and 2) whole or dissected tissue types from each of the three sampling sites discriminated separately was undertaken using a LDA score (log_10_) cut-off of 2.5 with *microbiomeMarker* (100). The V4 16S rDNA sequences from the entire bacterial community were next aligned with a staggered alignment using *DECIPHER* (2.14.0) using the profile-to-profile method (99). Using this alignment, a phylogenetic tree was constructed for the entire bacterial community with *Phangorn* (v2.5.5) (98) using the neighbor-joining tree estimation function (101) with the generalized time-reversible model (102) (including an optimized proportion of variable size and gamma rate parameters) and stochastic tree rearrangement.

To evaluate patterns in the bacterial communities detected among whole or dissected individual *I. scapularis*, distance measures between the V4 16S rDNA libraries were calculated using phylogeny-aware weighted and unweighted UniFrac distances (103), ordinated by PCoA using *phyloseq* and *ape* (v5.4-1) (104) and labeled by sample location and tissue type. For unweighted UniFrac ordinations, convex hull Gaussian ellipses were estimated using the Khachiyan algorithm (105) as implemented in the R package *ggforce* (v0.3.3) (106). A Venn diagram depicting common or unique ASVs detected in whole or dissected tick-associated libraries was generated with *VennDiagram* (v1.6.20) (107).

A permutation test of difference in multivariate homogeneity of group dispersions (variances) between tissue and whole tick libraries based on the unweighted UniFrac PCoA (see above) was implemented using *vegan* (v2.5-7). Non-parametric Wilcoxon rank sum tests (108) were used to identify significant differences between mean unweighted UniFrac distances calculated from within and across libraries representing whole ticks and dissected tissues. To investigate how rare taxa influenced the results, the statistical analysis was repeated with the 500 most abundant ASVs (rather than the agglomerated dataset described above) following normalization using a total-sum scaling (counts from each ASV divided by total library size).

### Metatranscriptome analysis

Paired-end sequence data were trimmed for quality and adapter removal of Illumina-specific adapter sequences (Supplemental Table S6) using *Flexbar* (v3.0.3) with trim-end mode ‘ANY’, minimum adapter overlap of 5, the maximum uncalled base of 1, and a minimum quality threshold of 20 (109). The metatranscriptome libraries from the four ticks were screened for mostly (92 – 96 %) *I. scapularis* reads from the genome assembly (phased diploid assembly from ISE6 cell line (Accession: GCF_00289285.2) (110) and PhiX (Accession: MN385565.1) using the splice-aware global aligner *BBMap* (v38.75) (111). The maximum indel size was increased to 200,000 against the host to accommodate large introns in the tick genome (112). The metatranscriptome was also filtered for residual amounts of prokaryotic and eukaryotic rDNA (∼1 – 4 %) using the *Sortmerna* (v4.0.0) package (113). The remaining ∼ 2 – 3 % resultant filtered sequence data were carried into the *de novo* assembly.

The metatranscriptome was assembled *de novo* using *Trinity* (v2.8.4) in default mode (114) and then clustered at 90 % sequence identity threshold using *CD-HIT-EST* (v4.6.8) (115). Secondary clustering generated 6,868 assembled transcripts (N50 = 2,063 bp; L50 = 320 bp) but left a high proportion (82.8 %) of unassembled sequences. Count tables were generated by aligning the filtered sequence data against the reference transcripts using *Salmon* (v0.14.1) (116). Successfully aligned sequences represented 29.1 % of all paired-end reads. Prior to normalization, the raw count data were filtered for low abundance transcripts (i.e., fewer than five aligned reads in at least one sample). The count data were then normalized with the upper quartile scaling approach and transformed using the *voom* function implemented in *edgeR* (117) and *limma* (118, 119), respectively. The log_2_ counts-per-million (CPM) expression values were visualized with *ggplot2* (95).

For taxonomic assignment to each of the reference transcripts, we used *BLASTx* (120) to search against GenBank’s non-redundant (nr) protein database (97). Similarly, we also used *BLASTn* (120) to search against GenBank’s nucleotide (nt) database (97) and the *R. buchneri* genome assembly REISMNv1 (Accession: GCF_000696365.1) (121). An e-value retention threshold of 1e^-5^ was used for all *BLAST*-based searches. Taxonomy could be assigned to 2,206 (32.1 %) reference transcripts by priority using matches from the *R. buchneri* genome with the lowest e-value, over matches against any residual *I. scapularis* non-redundant proteins, then followed by the lowest e-value from either the nr protein or nucleotide reference databases. Taxonomic information was assigned based on the blast-generated taxonIDs using *TaxonKit* (v0.5.0) (122) and the final metatranscriptome was filtered for transcripts annotated as unwanted Phyla, including *Arthropoda*, *Brachiopoda*, *Chordata*, *Cnidaria*, *Echinodermata*, *Mollusca*, *Priapulida*, *Streptophyta*, and *Tardigrada*.

## Data availability statement

Accession numbers SRR17194086 & SRR17194087 have been deposited by the authors into GenBank’s Sequence Read Archive (SRA) representing the raw sequence data analyzed in this study. The authors have made these data available without restriction.

The authors have also made available, without restriction, the details related to the programs and algorithms used to analyze these sequencing data, including suitable documentation regarding their use, as a reproducible analysis pipeline through GitHub. Please see https://github.com/ damselflywingz/tick_microbiome (archived version https://doi.org/10.5061/dryad.fqz612jw9) and the above methods and materials for the relevant details. This analysis pipeline also includes the computer code created to interpret data and generate the results presented in this study.

## Acknowledgments

The authors would like to acknowledge C. Brdar and E. Barkley of Ontario Parks, and S. Knapton of the Cataraqui Region Conservation Authority for authorizing the collection of tick samples used in this study. The authors are grateful to K. Ding and A. Siew of Queen’s University Biology Department, who performed the tick dissections and extractions. The authors are also grateful to Z. Sun and G. McLeod of Queen’s University Biology Department, who carried out the library preparations. The authors appreciate Dr. Sima Afsharnezhad and Logan Wisteard for debugging the bioinformatic pipeline. The authors acknowledge Dr. Samir Patel provided feedback on an early version of the manuscript.

## Author Contributions

**Amber R. Paulson:** Bioinformatics, formal analysis, data curation, writing – original draft, visualization. **David Huang:** Bioinformatics, validation, writing – review & editing **Stephen C. Lougheed:** Conceptualization, methodology, investigation, writing – review & editing. **Robert I. Colautti:** Conceptualization, methodology, validation, investigation, resources, writing – review & editing, supervision, project administration, funding acquisition.

## Legend for Supplemental Material

Supplemental Table S1: Core microbiome-associated ASVs detected in this study.

Supplemental Table S2: *Rickettsia*-associated ASVs detected in this study.

Supplemental Figure S1: Total abundance of core bacterial amplicon sequence variants associated with *I. scapularis*.

Supplemental Table S3: Raw count abundance of *Rickettsia*-associated Amplicon Sequence Variants (ASVs) from batch replicate 16S rRNA libraries.

Supplementary Figure S2: Relative abundance of core amplicon sequence variants associated with *I. scapularis* batch PCR replicate libraries.

Supplemental Table S4: Contaminating ASVs detected using decontam R package.

Supplemental Figure S3: Rarefaction curves for V4 16S rDNA sequencing libraries representing the bacterial community associated with whole and dissected I. scapularis.

Supplemental Table S5: *Babesia* apicoplast sequence detected (“ASV7”).

Supplemental Figure S4: Phylogenetic reconstruction for partial apicoplast sequence of *Babesia* spp. detected in the microbiome of *I. scapularis*.

Supplemental Figure S5: Phylogenetic reconstruction of V4 16S rDNA sequences from *A*. *phagocytophilum* target ASV6 detected in the microbiome of *I. scapularis* in this study.

Supplemental Figure S6: Phylogenetic reconstruction of V4 16S rRNA DNA sequences from *Borrelia* targets ASV2 and ASV2 detected in the microbiome of *I. scapularis*.

Supplemental Figure S7: LEfSe analysis of tick microbiome for discriminating ASVs.

Supplemental Figure S9: Pairwise comparison (Comp) of unweighted UniFrac distances between tick-associated bacterial communities.

Supplemental Figure S10: Repeated analysis with the top 500 most abundant amplicon sequence variants.

Supplemental Table S6: illumina_nextera.fa.

