## Supplementary materials for "Multi-omics analysis identifies symbionts and pathogens of blacklegged ticks (*Ixodes scapularis*) from a Lyme disease hotspot in southeastern Ontario, Canada"

Pre-print version

Note: See https://github.com/damselflywingz/tick_microbiome/ for codes and details related to the reproducible analysis pipeline for this study (archived version https://doi.org/10.5061/dryad.fqz612jw9).

Pre-print version

Supplemental Table S1: Core microbiome-associated ASVs detected in this study.

20220502_core_job
11.fasta

*>Rickettsia sp.*
*CCTGTTTGCTCCCCACGCTTTCGTGCATTAGCGTCAGTTGTAGCCCAGATGACCGCCTTC*
*GCCACCGGTGTTCCTCCTAATATCTAAGAATTTCACCTCTACACTAGGAATTCCATCATCCC*
*CTACTACACTCTAGATTAGTAGTTTTGAAAGCAATTCCGAGGTTAAGCCCCGGGCTTTCAC*
*TTCCAACTTACTAAACCGCCTACGCACTCTTTACGCCCAGTAATTCCGAACAACGCTAGCC*
*CCCTCCGTCT*
*>Borrelia miyamotoi*
*CCTGTTTGCTCCCTACGCTTTCGTGACTCAGCGTCAGTCTTGACCTAGAAGTTCGCCTTCG*
*CCTCTGGTATTCTTCCTGATATCAACAGATTCCACCCTTACACCAGGAATTCTAACTTCCCC*
*TATCAGACTCTAGTCATGCAGTTTCCAGCATAGTTCCACAGTTGAGCTGTGGTATTTTACAC*
*ATAGACTTGCATATCCGCCTACTCACCCTTTACGCCCAATGATCCCGAACAACGCTCGCCC*
*CTTACGTATT*
*>Borrelia sp.*
*CCTGTTTGCTCCCTACGCTTTCGTGACTCAGCGTCAGTCTTGACCCAGAAGTTCGCCTTCG*
*CCTCCGGTATTCTTTCTGATATCAACAGATTCCACCCTTACACCAGAAATTCTAACTTCCTC*
*TATCAGACTCTAGACATATAGTTTCCAACATAGGTCCACAGTTGAGCTGTGGTATTTTATGC*
*ATAGACTTATATATCCGCCTACTCACCCTTTACGCCCAATAATCCCGAACAACGCTCGCCC*
*CTTACGTAT*
*>Sphingomonas sp.*
*CCTGTTTGCTCCCCACGCTTTCGCACCTCAGCGTCAATACATGTCCAGTCAGCCGCCTTC*
*GCCACTGGTGTTCTTCCGAATATCTACGAATTTCACCTCTACACTCGGAATTCCACTGACCT*
*CTCCATGATTCAAGCGATGCAGTCTAAAAGGCAATTCCAGAGTTGAGCTCTGGGCTTTCAC*
*CTCTTACTTACAAAGCCGCCTACGTGCGCTTTACGCCCAGTAATTCCGAATAACGCTAGCT*
*CCCTCCGTA*
*>Pseudomonas sp.*
*CCTGTTTGCTCCCCACGCTTTCGCACCTCAGTGTCAGTATCAGTCCAGGTGGTCGCCTTC*
*GCCACTGGTGTTCCTTCCTATATCTACGCATTTCACCGCTACACAGGAAATTCCACCACCC*
*TCTACCGTACTCTAGCTTGCCAGTTTTGGATGCAGTTCCCAGGTTGAGCCCGGGGCTTTCA*
*CATCCAACTTAACAAACCACCTACGCGCGCTTTACGCCCAGTAATTCCGATTAACGCTTGC*
*ACCCTCTGTATT*
*>Anaplasma phagocytophilum*
*CCTGTTTGCTCCCCACGCTTTCGCACCTCAGCGTCAGTACCGGACCAGATAGCCGCCTTC*
*GCCACTGGTGTTCCTCCTAATATCTACGAATTTCACCTCTACACTAGGAATTCCGCTATCCT*
*CTTCCGGACTCTAGTCTGGCAGTATTAAAAGCAGCTCCAGGGTTAAGCCCTGGCATTTCAC*
*CTTTAACTTACCGAACCGCCTACATGCCCTTTACGCCCAATAATTCCGAACAACGCTTGCC*
*CCCTCCGTATTACC*
*>Frondihabitans sp.*

Pre-print version

*CCTGTTCGCTCCCCATGCTTTCGCTCCTCAGCGTCAGTTACGGCCCAGAGATCTGCCTTC*
*GCCATCGGTGTTCCTCCTGATATCTGCGCATTCCACCGCTACACCAGGAATTCCAATCTCC*
*CCTACCGCACTCTAGTCTGCCCGTACCCACTGCAGGCTGGGGTTGAGCCCCCAGATTTC*
*ACAGCAGACGCGACAAACCGCCTACGAGCTCTTTACGCCCAATAATTCCGGACAACGCTT*
*GCACCCTACGTATTAC*
*>Mycobacterium sp.*
*CCTGTTCGCTCCCCACGCTTTCGCTCCTCAGCGTCAGTTACTGCCCAGAGACCCGCCTTC*
*GCCACCGGTGTTCCTCCTGATATCTGCGCATTCCACCGCTACACCAGGAATTCCAGTCTC*
*CCCTGCAGTACTCCAGTCTGCCCGTATCGCCCGCACGCCCACAGTTGAGCTGTGAGTTTT*
*CACGAACAACGCGACAAACCACCTACGAGCTCTTTACGCCCAGTAATTCCGGACAACGCT*
*CGGACCCTACGTA*
*>Curtobacterium sp.*
*CCTGTTCGCTCCCCATGCTTTCGCTCCTCAGCGTCAGTTACGGCCCAGAGATCTGCCTTC*
*GCCATCGGTGTTCCTCCTGATATCTGCGCATTCCACCGCTACACCAGGAATTCCAATCTCC*
*CCTACCGCACTCTAGTCTGCCCGTACCCACTGCAAGCCCGAGGTTGAGCCTCGGGATTTC*
*ACAGCAGACGCGACAAACCGCCTACGAGCTCTTTACGCCCAATAATTCCGGACAACGCTT*
*GCACCCTACGTATT*
*>Sphingomonas sp._1*
*CCTGTTTGCTCCCCACGCTTTCGCACCTCAGCGTCAATACCAGTCCAGTGAGCCGCCTTC*
*GCCACTGGTGTTCTTCCGAATATCTACGAATTTCACCTCTACACTCGGAATTCCACTCACCT*
*CTCCTGGATTCAAGCGATGCAGTCTTAAAGGCAATTCTGGAGTTGAGCTCCAGGCTTTCAC*
*CTCTAACTTACAAAGCCGCCTACGTGCGCTTTACGCCCAGTAATTCCGAATAACGCTAGCT*
*CCCTCCGTA*
*>Shinella sp.*
*CCTGTTTGCTCCCCACGCTTTCGCACCTCAGCGTCAGTAATGGACCAGTAAGCCGCCTTC*
*GCCACTGGTGTTCCTCCGAATATCTACGAATTTCACCTCTACACTCGGAATTCCACTTACCT*
*CTTCCATACTCTAGGTACCCAGTATCAAAGGCAGTTCCAGAGTTGAGCTCTGGGATTTCAC*
*CCCTGACTTAAATACCCGCCTACGTGCGCTTTACGCCCAGTAATTCCGAACAACGCTAGCC*
*CCCTTCGTATT*
*>Williamsia sp.*
*CCTGTTCGCTACCCACGCTTTCGCTCCTCAGCGTCAGTTACTACCCAGAGACCCGCCTTC*
*GCCACCGGTGTTCCTCCTGATATCTGCGCATTTCACCGCTACACCAGGAATTCCAGTCTCC*
*CCTGTAGTACTCAAGTCTGCCCGTATCGCCCGCACGCTTGATGTTAAGCATCAAGATTTCA*
*CGAACGACGCGACAAACCGCCTACGAGCTCTTTACGCCCAGTAATTCCGGACAACGCTCG*
*CACCCTACGTATT*
*>Methylobacterium sp.*
*CCTGTTTGCTCCCCACGCTTTCGCGCCTCAGCGTCAGTGTCGGTCCAGTTGGCCGCCTTC*
*GCCACCGGTGTTCTTGCGAATATCTACGAATTTCACCTCTACACTCGCAGTTCCACCAACC*
*TCTACCGAACTCAAGCCATCCAGTATCGAAGGCAATTCTGTGGTTGAGCCACAGGCTTTCA*
*CCCCCGACTTAAATGGCCGCCTACGCGCCCTTTACGCCCAGTGATTCCGAGCAACGCTAG*
*CCCCCTTCGTATTAC*
*>Acidovorax sp.*

Pre-print version

CCTGTTTGCTCCCCACGCTTTCGTGCATGAGCGTCAGTACAGGTCCAGGGGATTGCCTTC
GCCATCGGTGTTCCTCCGCATATCTACGCATTTCACTGCTACACGCGGAATTCCATCCCCC
TCTACCGTACTCTAGCTATACAGTCACAAATGCAGTTCCCAGGTTGAGCCCGGGGATTTCA
CATCTGTCTTATATAACCGCCTGCGCACGCTTTACGCCCAGTAATTCCGATTAACGCTTGC
ACCCTACGTATT
>Pedobacter sp.
CCTGTTCGATCCCCACGCTTTCGTGCCTCAGCGTCAATAGGACCATAGTAAGCTGCCTTC
GCAATCGGTGTTCTGTGACATATCTATGCATTTCACCGCTACTTGTCACATTCCGCCTACCT
CTAGTCCATTCAAGCCCATCAGTATCAAGGGCACTGCGATGGTTGAGCCACCGTCTTTCAC
CCCTGACTTAACAGGCCGCCTACGCACCCTTTAAACCCAATAAATCCGGATAACGCTTGGA
TCCTCCGTATT
>Methylobacterium sp._1
CCTGTTTGCTCCCCACGCTTTCGCGCCTCAGCGTCAGTAATGGTCCAGTTGGCCGCCTTC
GCCACCGGTGTTCTTGCGAATATCTACGAATTTCACCTCTACACTCGCAGTTCCACCAACC
TCTACCATACTCAAGCGTCCCAGTATCGAAGGCCATTCTGTGGTTGAGCCACAGGCTTTCA
CCCCCGACTTAAAACGCCGCCTACGCGCCCTTTACGCCCAGTGATTCCGAGCAACGCTAG
CCCCCTTCGTATT
>Methylobacterium sp._2
CCTGTTTGCTCCCCACGCTTTCGCGCCTCAGCGTCAGTGTTGGTCCAGTTGGCCGCCTTC
GCCACTGGTGTTCTTGCGAATATCTACGAATTTCACCTCTACACTCGCAGTTCCACCAACC
TCTACCAAACTCAAGCCAAACAGTATCGAAGGCAATTCTGTGGTTGAGCCACAGGCTTTCA
CCCCCGACTTGAATGGCCGCCTACGCGCCCTTTACGCCCAGTGATTCCGAGCAACGCTAG
CCCCCTTCGTATTAC
>Massilia sp.
CCTGTTTGCTCCCCACGCTTTCGTGCATGAGCGTCAATCTTGACCCAGGGGGCTGCCTTC
GCCATCGGTGTTCCTCCACATCTCTACGCATTTCACTGCTACACGTGGAATTCTACCCCCC
TCTGCCAGATTCAAGCCTTGCAGTCTTCATCGCAATTCCCAGGTTGAGCCCGGGGCTTTCA
CGACAAACTTACAAAACCGCCTGCGCACGCTTTACGCCCAGTAATTCCGATTAACGCTTGC
ACCCTACGTATTAC
>Comamonas sp.
CCTGTTTGCTCCCCACGCTTTCGTGCATGAGCGTCAGTGCAGGCCCAGGGGATTGCCTTC
GCCATCGGTGTTCCTCCGCATATCTACGCATTTCACTGCTACACGCGGAATTCCATCCCCC
TCTGCCGCACTCTAGCTTTGCAGTCACAATGGCAGTTCCCAGGTTGAGCCCGGGGATTTC
ACCACTGTCTTACAAAACCGCCTGCGCACGCTTTACGCCCAGTAATTCCGATTAACGCTTG
CACCCTACGTATT
>Curvibacter sp.
CCTGTTTGCTCCCCACGCTTTCGTGCATGAGCGTCAGTACAGGCCCAGGGGATTGCCTTC
GCCATCGGTGTTCCTCCGCATATCTACGCATTTCACTGCTACACGCGGAATTCCATCCCCC
TCTGCCGTACTCTAGCTATGCAGTCACAAATGCAGTTCCCAGGTTGAGCCCGGGGATTTC
ACATCTGTCTTACATAACCGCCTGCGCACGCTTTACGCCCAGTAATTCCGATTAACGCTCG
CACCCTACGTATT
>Methyloversatilis universalis

Pre-print version

*CCTGTTTGCTCCCCACGCTTTCGTGCATGAGCGTCAGTATTGGCCCAGGGGGCTGCCTTC GCCATCGGTGTTCCTCCACATCTCTACGCATTTCACTGCTACACGTGGAATTCCACCCCCC TCTGCCATACTCTAGCCGTGCAGTCACAAGCGCAGTTCCCAGGTTAAGCCCGGGGATTTC ACACCTGTCTTACACAACCGCCTGCGCACGCTTTACGCCCAGTAATTCCGATTAACGCTCG CACCCTACGTATT*
*>Hymenobacter sp.*
*CCTGTTCGCTCCCCACGCTGTCGTGCCTCAGCGTCAGTAACAGCCTAGTCAGCTGCCTTC GCAATCGGGGTTCTGGACTGTATCTATGCATTTCACCGCTACTCAGTCCATTCCGCCAACC TCGTCTGTACTCAAGCCTCACAGTATCCAGGGCAGTTCCGTTGTTGAGCAACGGGCTTTCA CCCCGGACTTATAAGGCCGCCTACGCACCCTTTAAACCCAATAAATCCGGACAACGCTCG CACCCTCCGTATTACCG*
*>Hymenobacter sp._1*
*CCTGTTCGCTCCCCACGCTGTCGTGCCTCAGCGTCAGTAACAGCCTAGTCAGCTGCCTTC GCAATCGGGGTTCTGGACTGTATCTATGCATTTCACCGCTACTCAGTCCATTCCGCCAACC TCGTCTGTACTCAAGCCCGTCAGTATCCAGGGCAGTTCCGTTGTTGAGCAACGGGCTTTC ACCCCGGACTTAACGGGCCGCCTACGCACCCTTTAAACCCAATAAATCCGGACAACGCTC GCACCCTCCGTATTACCG*
*>Neobacillus sp.*
*CCTGTTTGCTCCCCACGCTTTCGCGCCTCAGCGTCAGTTACAGACCAGAAAGCCGCCTTC GCCACTGGTGTTCCTCCACATCTCTACGCATTTCACCGCTACACGTGGAATTCCGCTTTCC TCTTCTGTACTCAAGTCCCCCAGTTTCCAATGACCCTCCACGGTTGAGCCGTGGGCTTTCA CATCAGACTTAAAGGACCGCCTGCGCGCGCTTTACGCCCAATAATTCCGGACAACGCTTG CCACCTACGTATTACC*
*>Sphingomonas sp._2*
*CCTGTTTGCTCCCCACGCTTTCGCACCTCAGCGTCAATACCAGTCCAGTGAGCCGCCTTC GCCACTGGTGTTCTTCCGAATATCTACGAATTTCACCTCTACACTCGGAATTCCACTCACCT CTCCTGGATTCAAGCGATGCAGTCTTAAAGGCTATTCCGGAGTTGAGCCCCGGGCTTTCA CCTCTAACTTACAAAGCCGCCTACGTGCGCTTTACGCCCAGTAATTCCGAACAACGCTAGC TCCCTCCGTATTACC*

Supplemental Table S2: *Rickettsia*-associated ASVs detected in this study.

20220407_Rickettsia _6ASVs.fasta

>Rickettsia sp.
CCTGTTTGCTCCCCACGCTTTCGTGCATTAGCGTCAGTTGTAGCCCAGATGACCGCCT TCGCCACCGGTGTTCCTCCTAATATCTAAGAATTTCACCTCTACACTAGGAATTCCAT CATCCCCTACTACACTCTAGATTAGTAGTTTTGAAAGCAATTCCGAGGTTAAGCCCC GGGCTTTCACTTCCAACTTACTAAACCGCCTACGCACTCTTTACGCCCAGTAATTCCG AACAACGCTAGCCCCCTCCGTCT
>Rickettsia sp._1

Pre-print version

CCTGGTTGCTCCCCACGCTTTCGTGCATTAGCGTCAGTTGTAGCCCAGATGACCGCC
TTCGCCACCGGTGTTCCTCCTAATATCTAAGAATTTCACCTCTACACTAGGAATTCCA
TCATCCCCTACTACACTCTAGATTAGTAGTTTTGAAAGCAATTCCGAGGTTAAGCCC
CGGGCTTTCACTTCCAACTTACTAAACCGCCTACGCACTCTTTACGCCCAGTAATTCC
GAACAACGCTAGCCCCCTCCGTCT
>Rickettsia sp._2
CCTGTTTGCTCCCCACGCTTTCGTGCATTAGCGTCAGTTGTAGCCCAGATGACCGCCT
TCGCCACCGGTGTTCCTCCTAATATCTAAGAATTTCACCTCTACACTAGGAATTCCAT
CATCCCCTACTACACTCTAGATTAGTAGTTTTGAAAGCAATTCCGAGGTTAAGCCCC
GGGCTTTCACTTCCAACTTACTAAACCGCCTACGCACTCTTTACGCCCAGTAATTCCG
AACAACGCTCGCCCCTTACGTATTACC
>Rickettsia sp._3
CCTGTTTGCTCCCCACGCTTTCGTGCATTAGCGTCAGTTGCAGCCCAGATGACCGCCT
TCGCCACCGGTGTTCCTCCTAATATCTAAGAATTTCACCTCTACACTAGGAATTCCAT
CATCCCCTACTACACTCTAGATTAGTAGTTTTGAAAGCAATTCCGAGGTTAAGCCCC
GGGCTTTCACATCCAACTTACTAAACCGCCTACGCACTCTTTACGCCCAGTAATTCC
GAACAACGCTAGCCCCCTCCGTCTTACC
>Rickettsia sp._4
CCTGTTTGCTCCCCACGCTTTCGCGCCTCAGCGTCAGTTGTAGCCCAGATGACCGCCT
TCGCCACCGGTGTTCCTCCTAATATCTAAGAATTTCACCTCTACACTAGGAATTCCAT
CATCCCCTACTACACTCTAGTCTAGCAGTTTTGAAAGCAATTCCGAGGTTAAGCCTC
GGGCTTTCACTTCCAACTTACTAGACCGCCTACGCGCTCTTTACGCCCAGTAATTCCG
AACAACGCTAGCCCCCTCCGTCTT
>Rickettsia sp._5
CCTGTTTGCTCCCCACGCTTTCGCACCTCAGCGTCAGTACCGGACCAGATAGCCGCC
TTCGCCACTGGTGTTCCTCCTAATATCTAAGAATTTCACCTCTACACTAGGAATTCCA
TCATCCCCTACTACACTCTAGATTAGTAGTTTTGAAAGCAATTCCGAGGTTAAGCCC
CGGGCTTTCACTTCCAACTTACTAAACCGCCTACGCACTCTTTACGCCCAGTAATTCC
GAACAACGCTAGCCCCCTCCGTCTTACC

Pre-print version

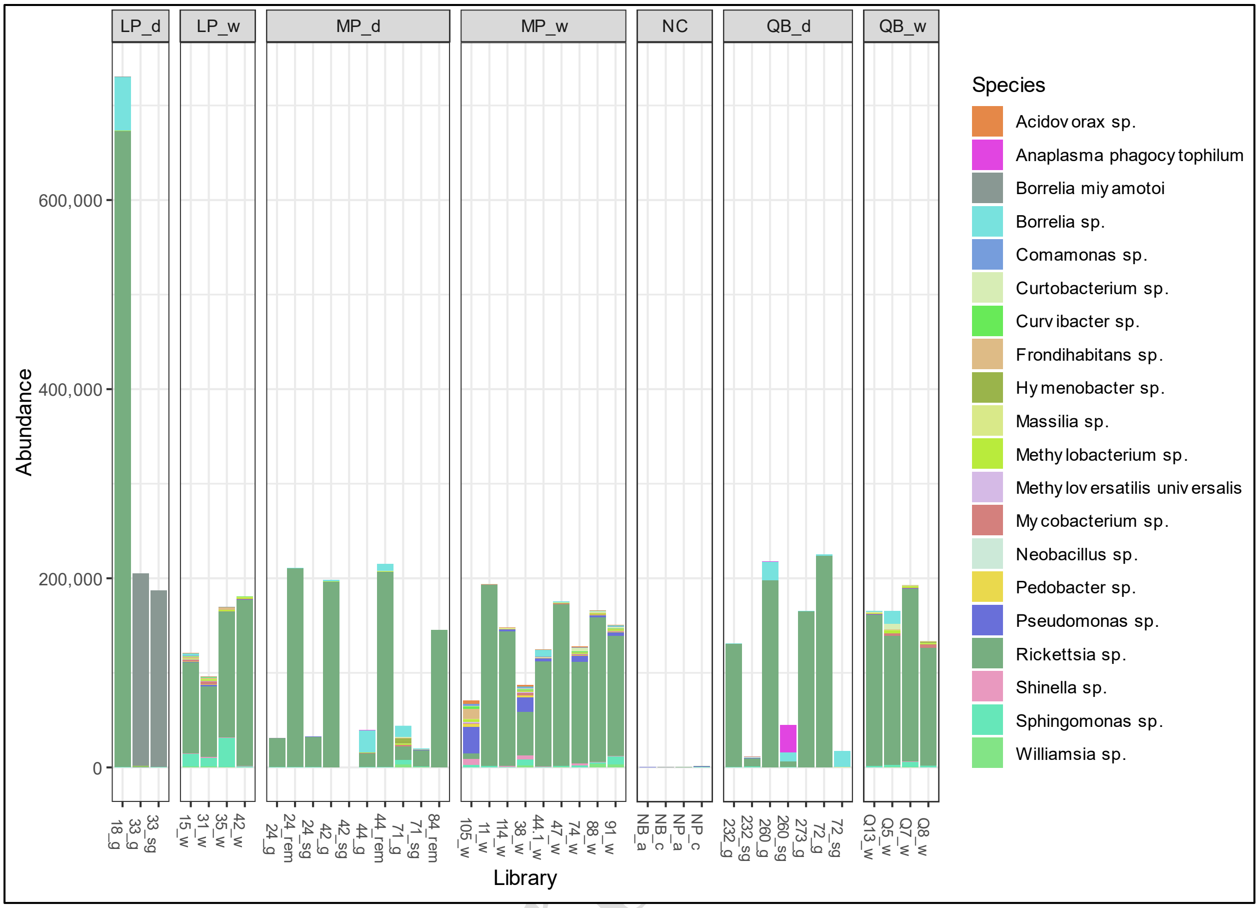

Supplemental Figure S1: Total abundance of core bacterial amplicon sequence variants (ASVs) associated with I. scapularis. Core bacterial community is represented by ASVs with a minimum relative abundance of 0.1 % in at least one V4 16S rDNA sequencing libraries. Sample types are indicated as salivary gland (sg), gut (g), remaining internal viscera (rem) or whole (w) in library names. Other abbreviations: Lemoine Point (LP), Murphy's Point (MP), Queen's University Biological Station (QB), negative control (NC), negative bead (NB), and negative PCR (NP).

Supplemental Table S3: Raw count abundance of *Rickettsia*-associated Amplicon Sequence Variants (ASVs) from batch replicate 16S rRNA libraries.

This file is available at: https://github.com/damselflywingz/tick_microbiome/blob/main/16S_microbiome/figs/Supplemental_Table_S3_.xls (archived version https://doi.org/10.5061/dryad.fqz612jw9)

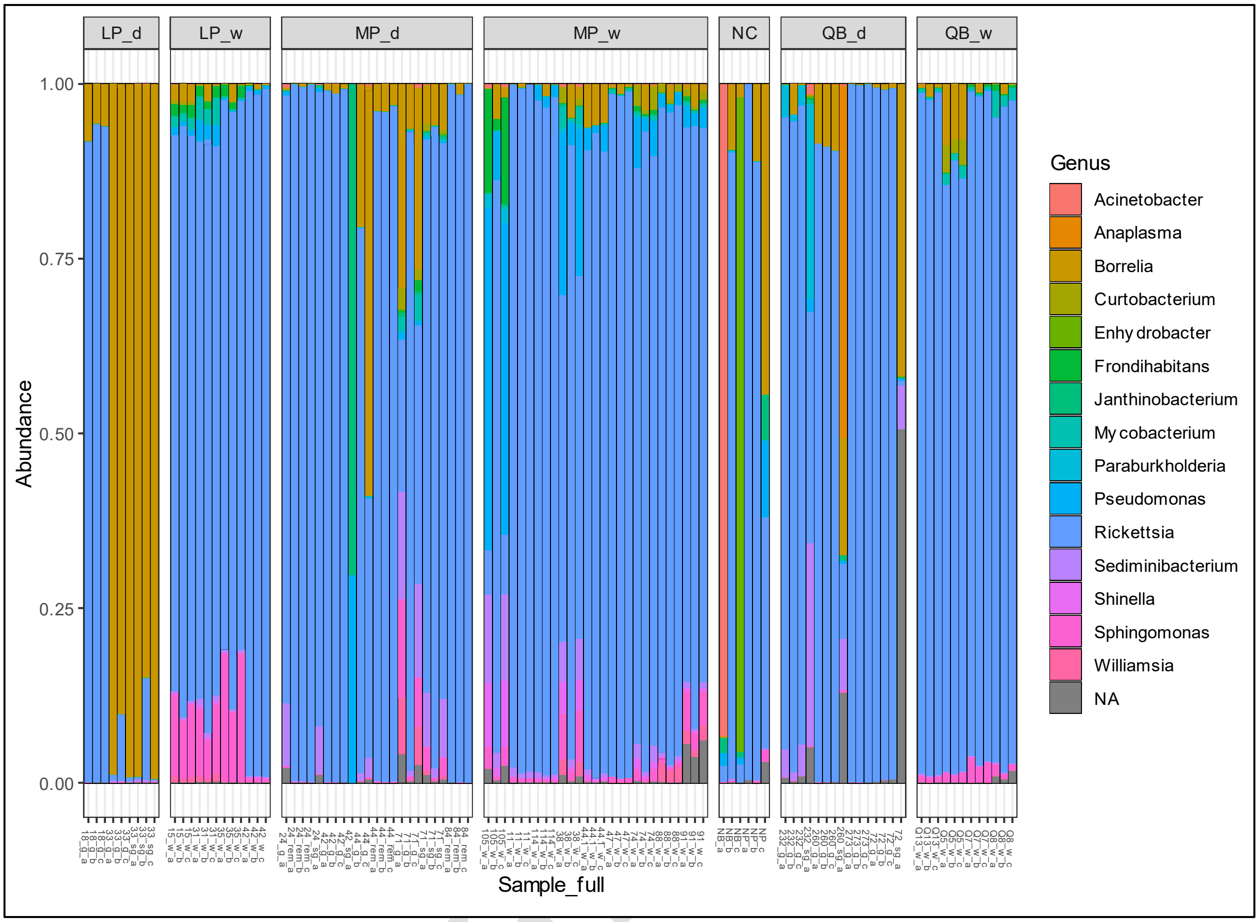

Supplementary Figure S2: Relative abundance of core amplicon sequence variants (ASVs) associated with *I. scapularis* batch PCR replicate libraries (denoted as “a”, “b” and “c” in the library names). Core bacterial community is represented by ASVs with a minimum relative abundance of 0.2 % in at least one V4 16S rDNA library. Sample types are indicated as salivary gland (sg), gut (g), remaining internal viscera (v) or whole (w) in library names. Control samples are indicated as negative bead (NB) and negative PCR (NP) in the library names. Other abbreviations: Lemoine Point (LP), Murphy’s Point (MP), Queen’s University Biological Station (QB), negative control (NC).

Supplemental Table S4: Contaminating ASVs detected using *decontam* R package.

This file is available at: https://github.com/damselflywingz/tick_microbiome/blob/main/16S_microbiome/figs/Supplemental_Table_S4_.xls. (archived version https://doi.org/10.5061/dryad.fqz612jw9)

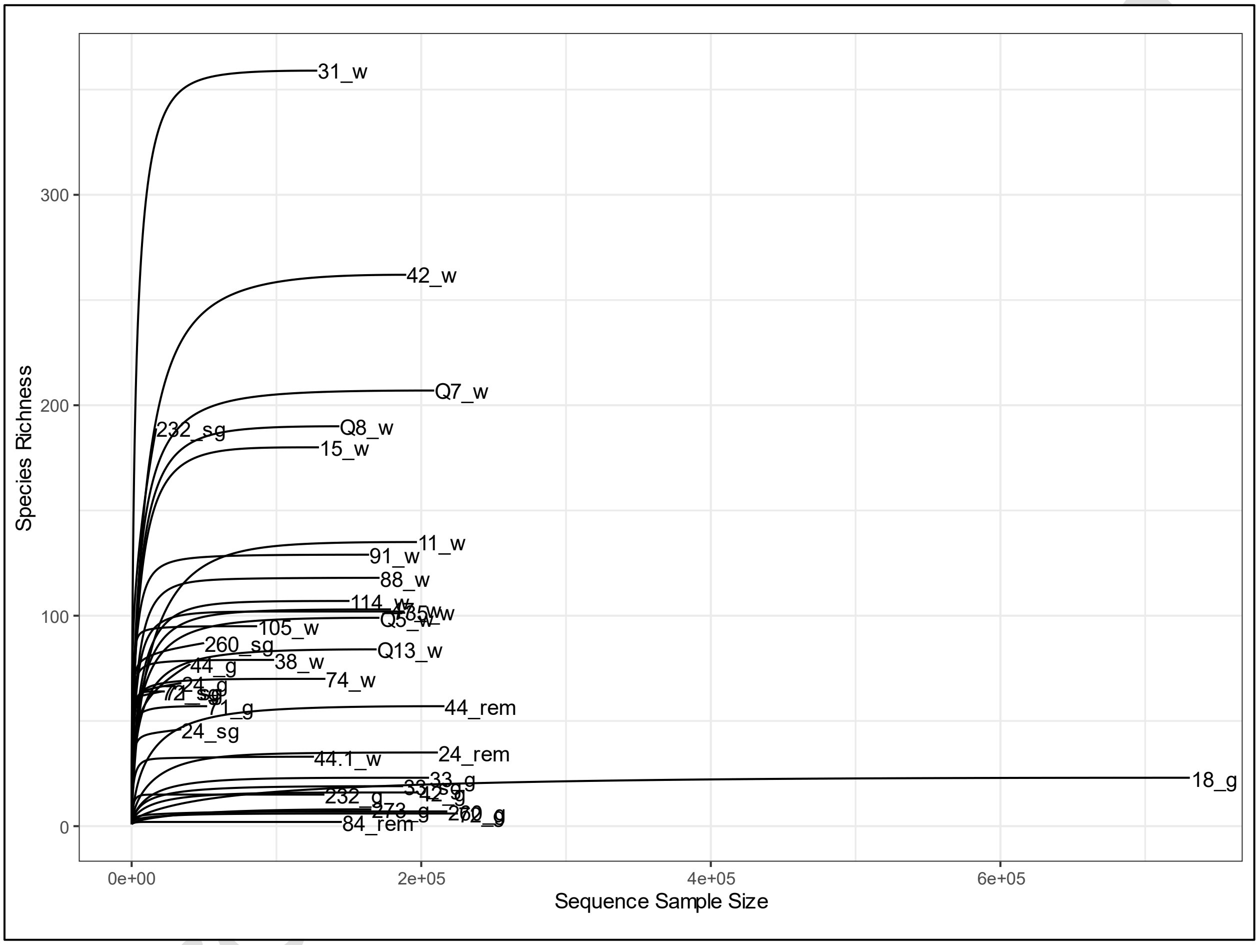

Supplemental Figure S3: Rarefaction curves for V4 16S rDNA sequencing libraries representing the bacterial community associated with whole and dissected *I. scapularis*. Species richness is the total number of amplicon sequence variants detected in each library. Sample types are indicated as salivary gland (sg), gut (g), remaining internal viscera (rem) or whole (w) in the library names. Control sample is indicated as negative bead (NB).

Supplemental Table S5: *Babesia* apicoplast sequence detected (“ASV7”).

20220407_ASV7.fasta

>Bacteria (kingdom)
CCCATTTGCTACTTTAGCTTTCATCTATCAATGTCAGATATAAACTAGTTTAATACTT
TCGTTTATGGCCTTCTTTAATGTATATAAATCATATTTCACCATTATTCAAAAAATTC
CTTAAACTTATTTTACTCTCAAGTTTATTAATATTAAATTTAATGTTTAAAAATATTA
TTTTTAAATTTATAATATAATAGAATAAACCATCTAAAGATGCTTTATGCCCAATAA
TTGTGAATAACGCTTATATCCTCTGTATTACC

Pre-print version

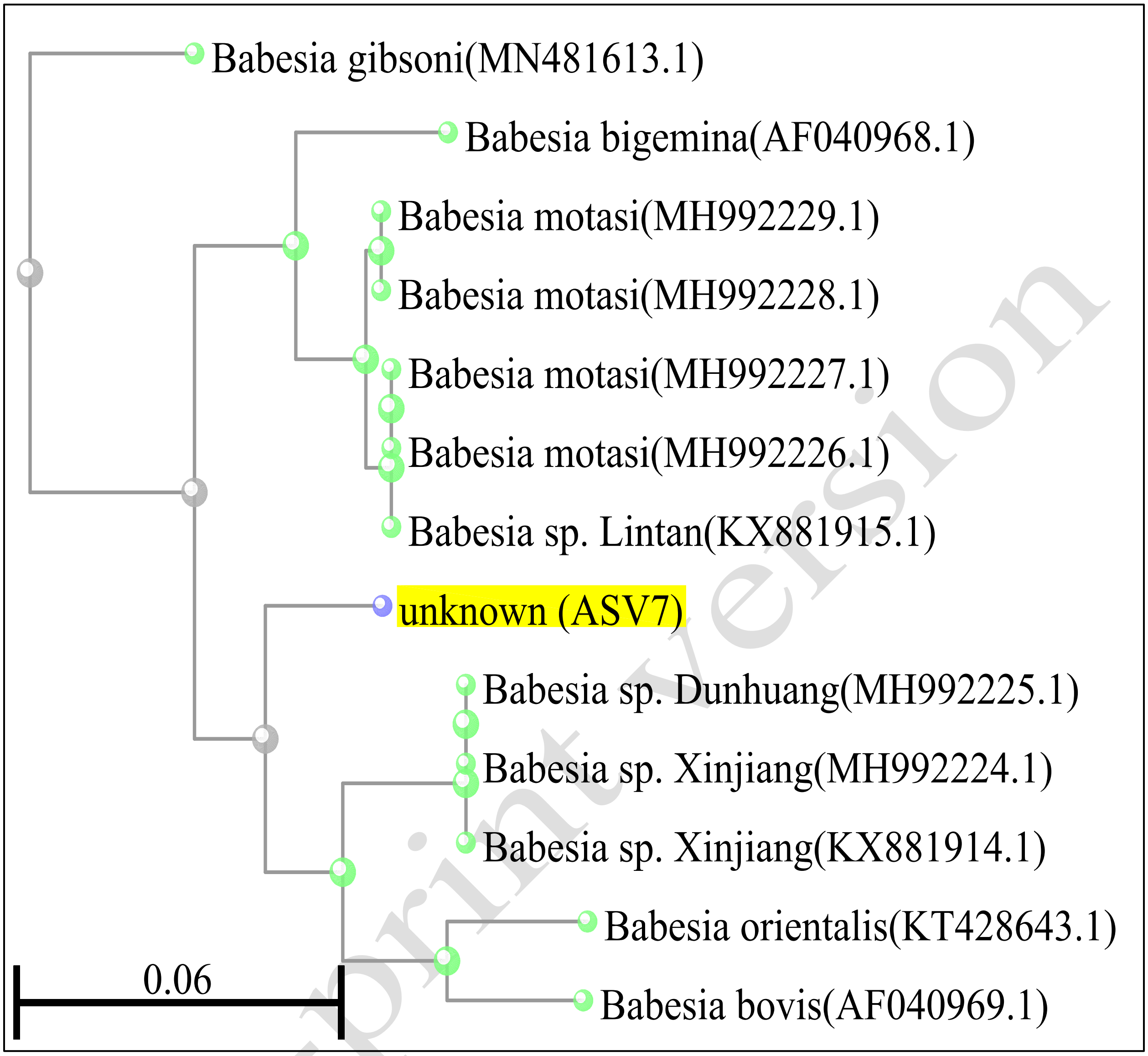

Supplemental Figure S4: Phylogenetic reconstruction for partial apicoplast sequence of *Babesia* spp. detected in the microbiome of *I. scapularis* by BLAST pairwise alignments, fast minimum evolution tree method with max sequence difference of 75 %.

Pre-print version

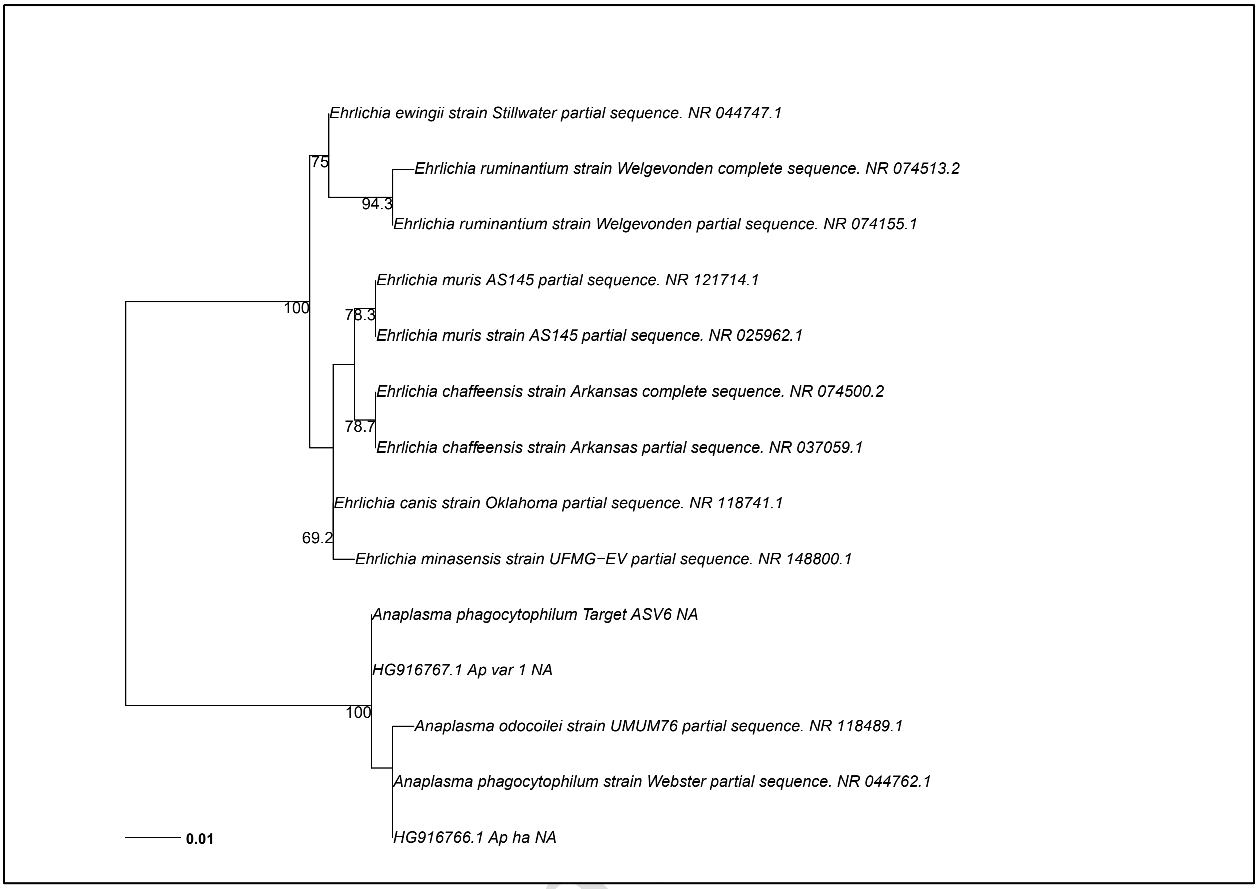

Supplemental Figure S5: Phylogenetic reconstruction of V4 16S rDNA sequences from *A. phagocytophilum* target ASV6 detected in the microbiome of *I. scapularis* in this study; 11 sequences from *A. phagocytophilum* strains and *Ehrlichia* spp. in the GenBank 16S rRNA reference database; and two *A. phagocytophilum* strains previously collected in Canada (Krakowetz *et al.* 2014). The bootstrap confidence is reported for internal nodes values greater than 65 % out of 1,000.

Pre-print

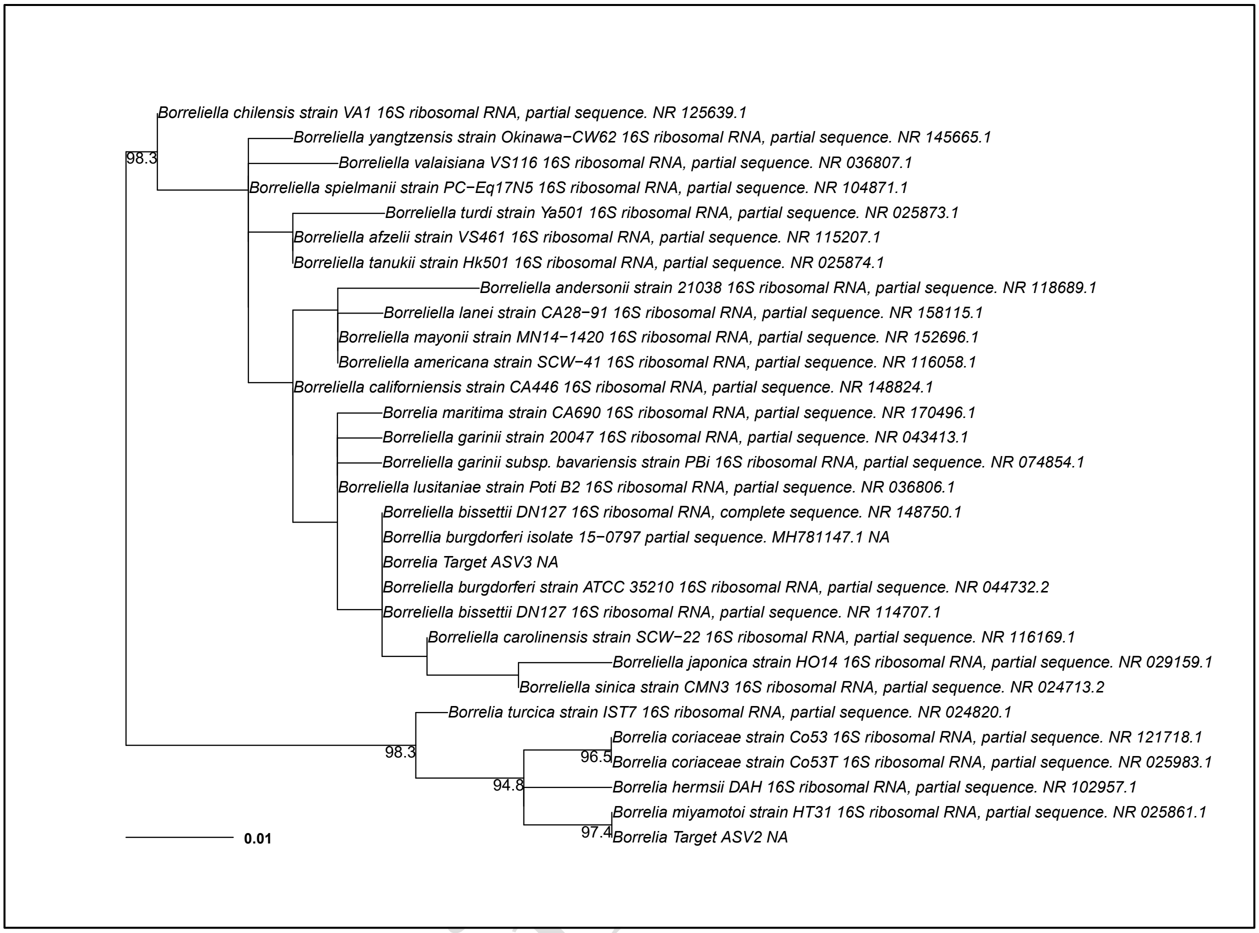

Supplemental Figure S6: Phylogenetic reconstruction of V4 16S rRNA DNA sequences from *Borrelia* targets ASV2 and ASV2 detected in the microbiome of *I. scapularis* , 27 sequences from *Borreliella/ Borrelia* spp. in the GenBank 16S rRNA reference database. Note that 16S rDNA gene sequence is not available for *Borreliella kurtenbachii*. The bootstrap confidence is reported for internal nodes values greater than 65 % out of 1,000.

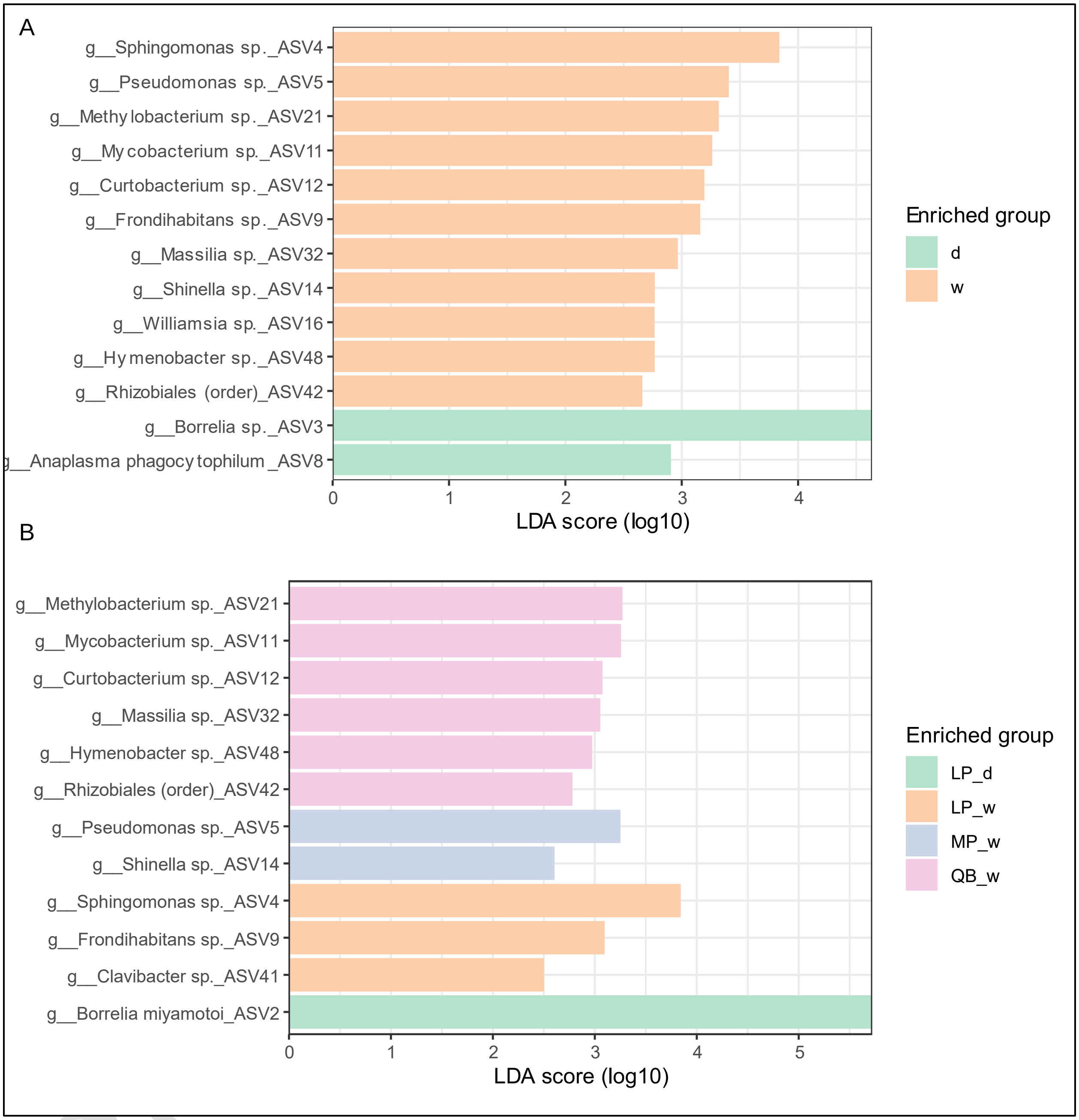

Supplemental Figure S7: LEfSe analysis of tick microbiome for discriminating ASVs associated with A) whole-tick (w), or dissected (d) salivary glands, guts, or remaining viscera samples across three sampling sites; and B) w or d tick samples at specific sites. The ASVs were agglomerated at the lowest taxonomic level; N = 419 (see methods). Sampling location abbreviations: Lemoine's Point (LP), Murphy's Point (MP) and Queen's University Biological Station (QB).

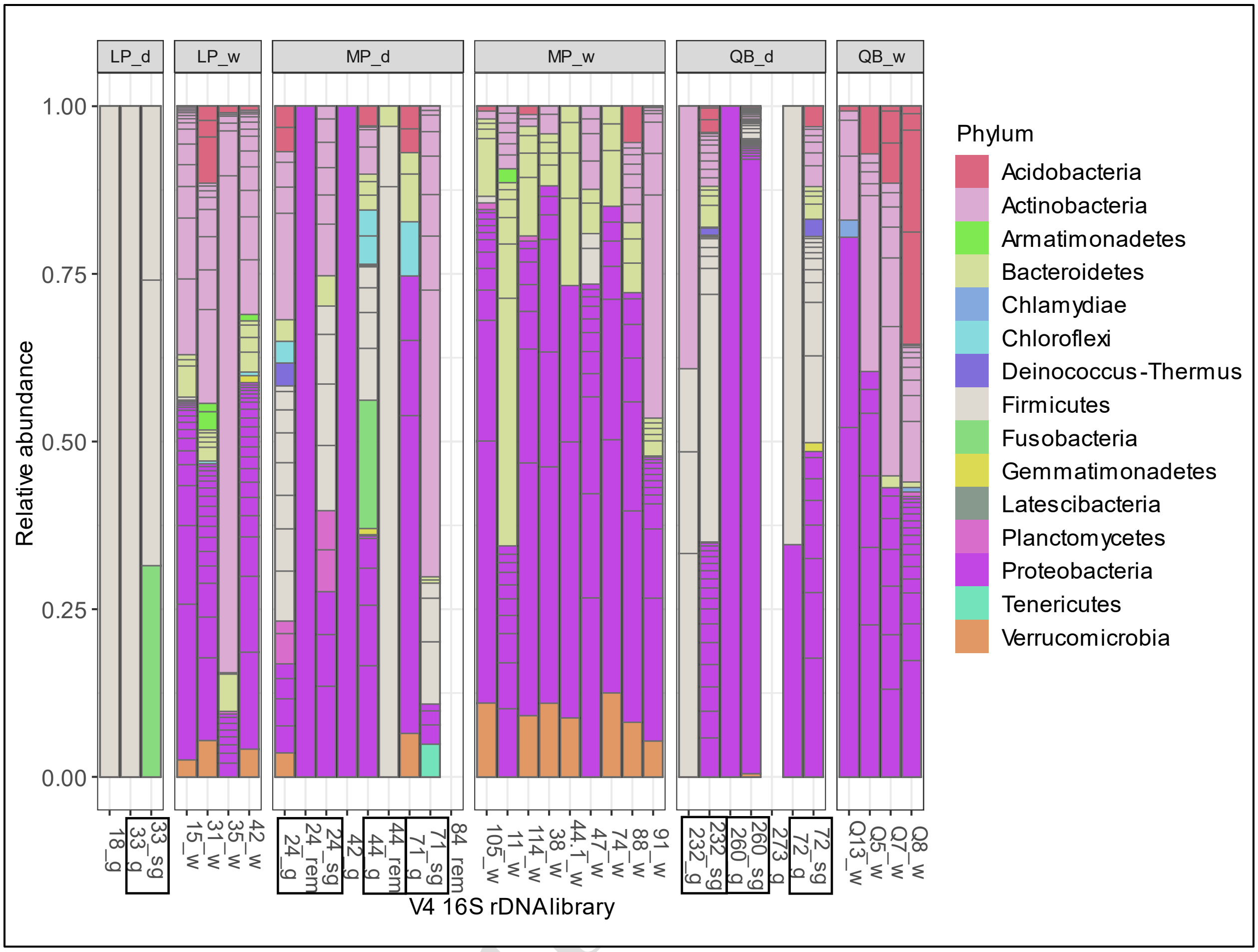

Supplemental Figure S8. Relative abundance of unique ASVs detected either from whole (W) *I. scapularis* or dissected (D) salivary glands (see Figure 3B in the main paper). Sample types are indicated in the library names as salivary gland (sg), gut (g), internal viscera (rem) or whole (w). Location abbreviations: Lemoine's Point (LP), Murphy's Point (MP) and Queen's University Biological Station (QB).

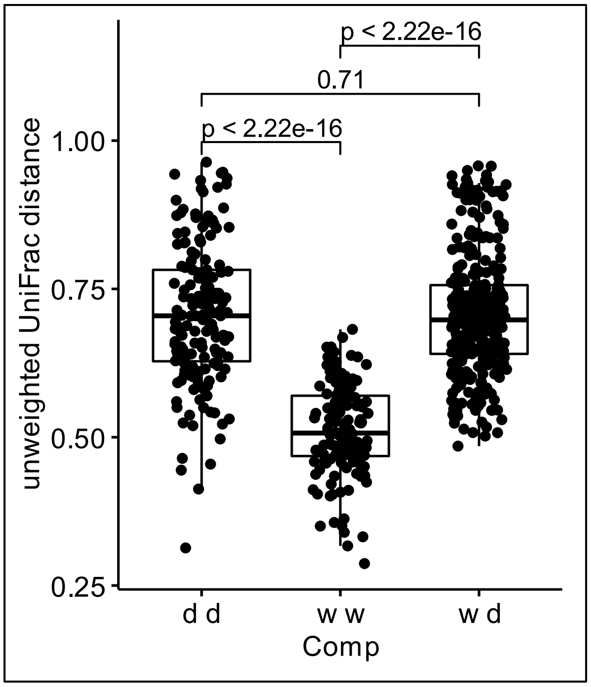

Supplemental Figure S9: Pairwise comparison (Comp) of unweighted UniFrac distances between tick-associated bacterial communities for within and between sample types of dissected (d) tissues or whole-body (w).

Pre-print version

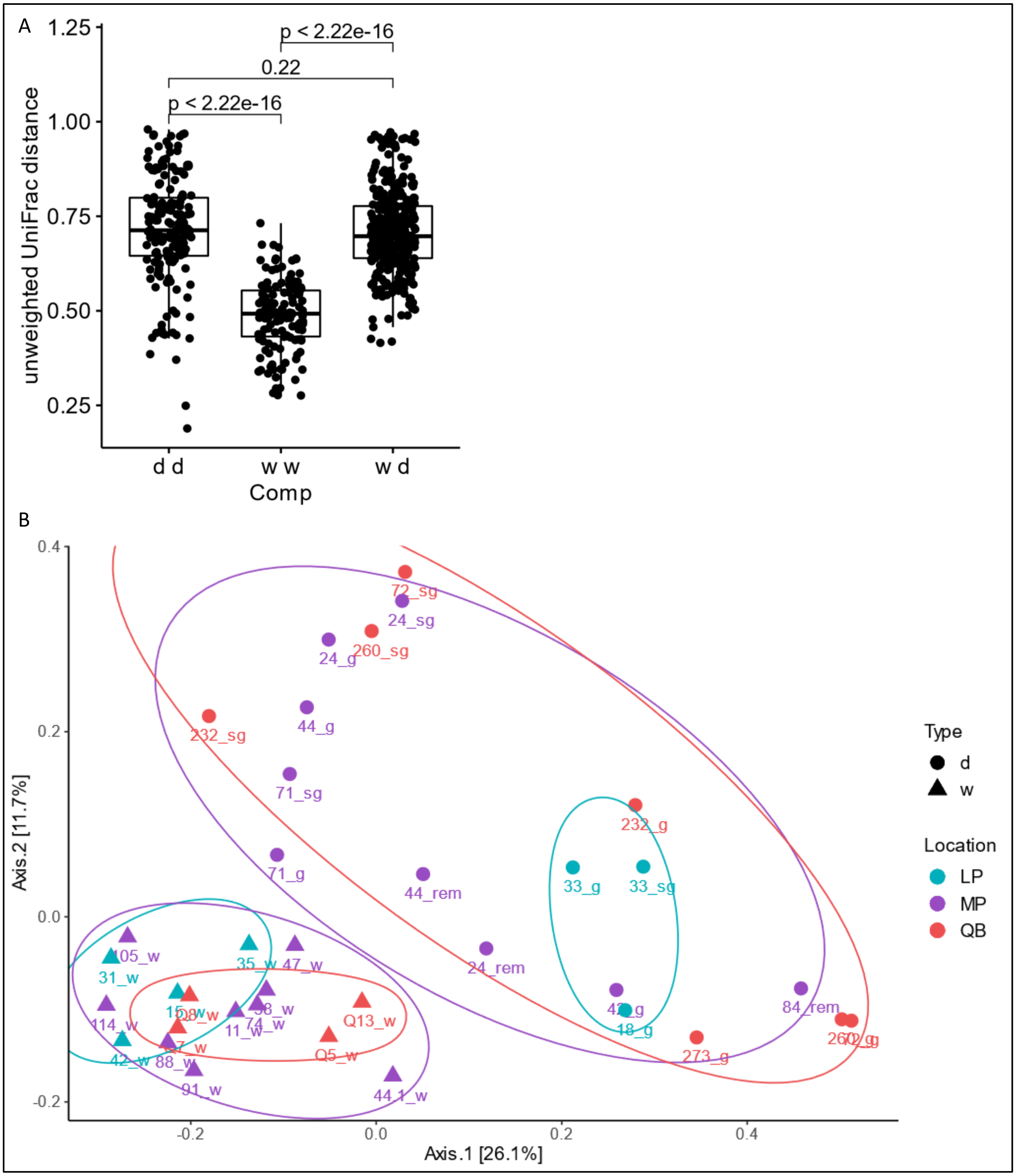
A
p < 2.22e-16
0.22
p < 2.22e-16
1.25
1.00
0.75
0.50
0.25
unweighted UniFrac distance
d d
w w
w d
Comp
B
0.4
0.2
0.0
-0.2
Axis.2 [11.7%]
-0.2
0.0
0.2
0.4
Axis.1 [26.1%]
72_sg
24_sg
24_g
260_sg
232_sg
44_g
71_sg
71_g
44_rem
24_rem
232_g
33_g
33_sg
18_g
84_rem
273_g
105_w
31_w
35_w
47_w
114_w
11_w
Q13_w
Q5_w
42_w
88_w
91_w
44.1_w
Type
d
w
Location
LP
MP
QB

Supplemental Figure S10: Repeated analysis with the top 500 most abundant amplicon sequence variants (ASVs). A) Pairwise comparison (Comp) of unweighted UniFrac distances between tick-associated bacterial communities for within and between sample types of dissected (d) tissues or whole-body (w). B) Principle co-ordinates analysis based on unweighted UniFrac distance metric ASVs detected from *I. scapularis* using V4 16S rDNA sequencing. The tissue types are indicated in the library names as either dissected (d): salivary gland (sg), gut (g), remaining (rem) internal viscera, or whole (w). Location abbreviations: Lemoine's Point (LP), Murphy's Point (MP) and Queen's University Biological Station (QB).

Supplemental Table S6 – illumina_nextera.fa

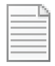

illumina_nextera.fa.txt

>Illumina_D5_adapter_i5a
ATGATACGGCGACCACCGAGATCTACAC
>Illumina_D5_adapter_i5b
ACACTCTTTCCCTACACGACGCTCTTCCGATCT
>Illumina_D5_adapter_i7a
GATCGGAAGAGCACACGTCTGAACTCCAGTCAC
>Illumina_D5_adapter_i7b
ATCTCGTATGCCGTCTTCTGCTTG
>Truseq_Universal_Adapter
AATGATACGGCGACCACCGAGATCTACACTCTTTCCCTACACGACGCTCTTCCGATC
T
>Truseq_Adapter_1a
GATCGGAAGAGCACACGTCTGAACTCCAGTCAC
>Truseq_Adapter_1b
ATCTCGTATGCCGTCTTCTGCTTG
>Illumina_D5_adapter_i5a Reversed:
GTGTAGATCTCGGTGGTCGCCGTATCAT
>Illumina_D5_adapter_i5b Reversed:
AGATCGGAAGAGCGTCGTGTAGGGAAAGAGTGT
>Illumina_D5_adapter_i7a Reversed:
GTGACTGGAGTTCAGACGTGTGCTCTTCCGATC
>Illumina_D5_adapter_i7b Reversed:
CAAGCAGAAGACGGCATACGAGAT
>Truseq_Universal_Adapter Reversed:
AGATCGGAAGAGCGTCGTGTAGGGAAAGAGTGTAGATCTCGGTGGTCGCCGTATCA
TT
>Truseq_Adapter_1a Reversed:
GTGACTGGAGTTCAGACGTGTGCTCTTCCGATC
>Truseq_Adapter_1b Reversed:
CAAGCAGAAGACGGCATACGAGAT

Pre-print version

>TruSeq_Universal_Adapter
AATGATACGGCGACCACCGAGATCTACACTCTTTCCCTACACGACGCTCTTCCGATC
T
>Nextera_XT
CTGTCTCTTATACACATCT
>Nextera_XT_Reverse
AGATGTGTATAAGAGACAG

Pre-print version
